# Deciphering single-cell heterogeneity and cellular ecosystem dynamics during prostate cancer progression

**DOI:** 10.1101/2024.12.18.629070

**Authors:** Faming Zhao, Jianming Zeng, Canping Chen, Xiaofan Zhao, Tingting Zhang, George V. Thomas, Rosalie C. Sears, Joshi J. Alumkal, Amy E. Moran, Gordon B. Mills, Peter S. Nelson, Zheng Xia

## Abstract

Prostate cancer (PC) progresses from benign epithelium through pre-malignant lesions, localized tumors, metastatic dissemination, and castration-resistant stages, with some cases exhibiting phenotype plasticity under therapeutic pressure. However, high-resolution insights into how cell phenotypes evolve across successive stages of PC remain limited. Here, we present the Prostate Cancer Cell Atlas (PCCAT) by integrating ∼710,000 single cells from 197 human samples covering a spectrum of tumor stages. This comprehensive analysis dissects the cellular landscape and characterizes key cell types and molecular features that associate with PC progression and prognosis. In malignant cells, we highlight a distinctive profile denoted by high Major Histocompatibility Complex (MHC) expression, low Androgen Receptor (AR) activity, and enhanced stemness programs associated with enzalutamide resistance. Moreover, we reveal several cell states strongly correlated with PC progression and adverse prognosis, including lineage plasticity-like malignant cells (LPCs), neuroendocrine tumor cells, pericytes, and matrix cancer-associated fibroblasts (mCAFs). Furthermore, we uncover shared cell states that underpin the immune suppressive tumor microenvironment in advanced PC, including activated regulatory T cells, exhausted CD8+ T cells, and SPP1-expressing macrophages. Lastly, we pinpoint a spatial niche composed of mCAFs and SPP1-expressing macrophages localized near the tumor boundary in aggressive PC, which correlates with poor prognosis. Overall, our work provides a valuable resource and offers deeper insights into the diverse cell states, dynamics, and functional characteristics involved in PC progression at single-cell resolution.

## Introduction

Prostate cancer (PC) is a highly heterogeneous disease, exhibiting diverse clinical, pathological, and molecular characteristics^1, 2^. Targeting Androgen Receptor (AR) signaling has proven effective in treating hormone sensitive PC (HSPC); however, the disease frequently progresses to a more aggressive subtype, known as castration-resistant PC (CRPC)^3^. Notably, approximately 17% of CRPC cases exhibit neuroendocrine features (CRPC-NE), with some advancing to a highly lethal and incurable subtype known as poorly differentiated neuroendocrine PC (NEPC)^1, 4, 5, 6^. The advent of next-generation sequencing has enabled a comprehensive characterization of the molecular landscape across various stages of PC progression, providing new insights into the pathobiology of the disease^3, 5, 7, 8, 9, 10, 11, 12, 13, 14, 15, 16, 17^. However, these studies have primarily relied on bulk tissue analyses which provide very limited views of cells comprising tumor microenvironments (TME) and their impact on PC progression.

Encouragingly, recent advances in single-cell RNA-seq (scRNA-seq) and spatial transcriptome RNA-seq (stRNA-seq) technologies have enabled the dissection of complex TMEs that accompany different stages of PC. Several reports have documented previously underestimated TME heterogeneity in early and advanced PC^18, 19, 20, 21, 22, 23, 24, 25, 26, 27, 28, 29, 30, 31, 32, 33, 34, 35, 36, 37^. Nevertheless, a major limitation of these assessments is the relatively small number of patient samples and cells analyzed in each study. Moreover, each provides only a snapshot of the overall picture, lacking comprehensive comparisons across multiple stages of PC progression. Additionally, the absence of genomic data as well as the lack of long-term follow-up information impedes a comprehensive dissection of the drivers of biological heterogeneity and its potential implications for therapy resistance and survival outcomes.

In the present work, we compiled publicly available scRNAseq datasets to address the aforementioned hurdles and created a comprehensive Prostate Cancer Cell Atlas (PCCAT) from 197 samples representing a range of benign and malignant prostate pathologies. This atlas offers a cohesive, high-resolution perspective of various PC stages, including normal healthy (N), benign prostate hyperplasia (BPH), normal adjacent to tumor (Adj), primary PC (Pri), invasive cribriform carcinoma and intraductal carcinoma (ICC/IDC), CRPC, metastatic HSPC (mHSPC) as well as metastatic CRPC (mCRPC) and NEPC. We performed an in-depth dissection of cell populations, uncovering the key cell types and molecular features associated with PC progression, treatment resistance, and prognosis. We also substantiated specific transcriptomic distinctions using spatial transcriptomics and bulk expression profiles. Finally, we developed an interactive web portal called PCCAT (https://pccat.net) to maximize broad access to this PC atlas and provide an automated mapping tool enabling the rapid cell type annotation of cells from PC, including subtypes of neoplastic epithelium and components of the TME.

## Results

### Single-cell transcriptome landscape across different stages of PC

Consensus definitions of cell types of PC, particularly those related to transitional cell states, based on single-cell transcriptomic data across studies, are currently lacking^38^. Therefore, we first developed PCCAT by compiling scRNA-seq data from 19 datasets comprising 197 samples from 143 individuals, providing a cohesive assessment of the cellular composition across stages of PC development and progression (see Methods for quality control and batch-effect correction; Figures 1A, S1, and S2; Table S1). Leveraging what is, to our knowledge, the largest and most comprehensive PC scRNA-seq atlas, we established a conceptual framework for harmonizing cell type nomenclature by building a three-level hierarchical cell identity reference, including major cell types (Level 1), refined cell subtypes (Level 2) and transitional cell states (Level 3) (Figure 1A), which enables a deeper understanding of the dynamic changes of cellular function under PC initiation and progression. As extended validation, we incorporated additional publicly-available scRNA-seq and stRNA-seq datasets, as well as bulk transcriptome cohorts comprising more than 3,000 samples (Figure 1B; Table S1).

**Figure 1.**
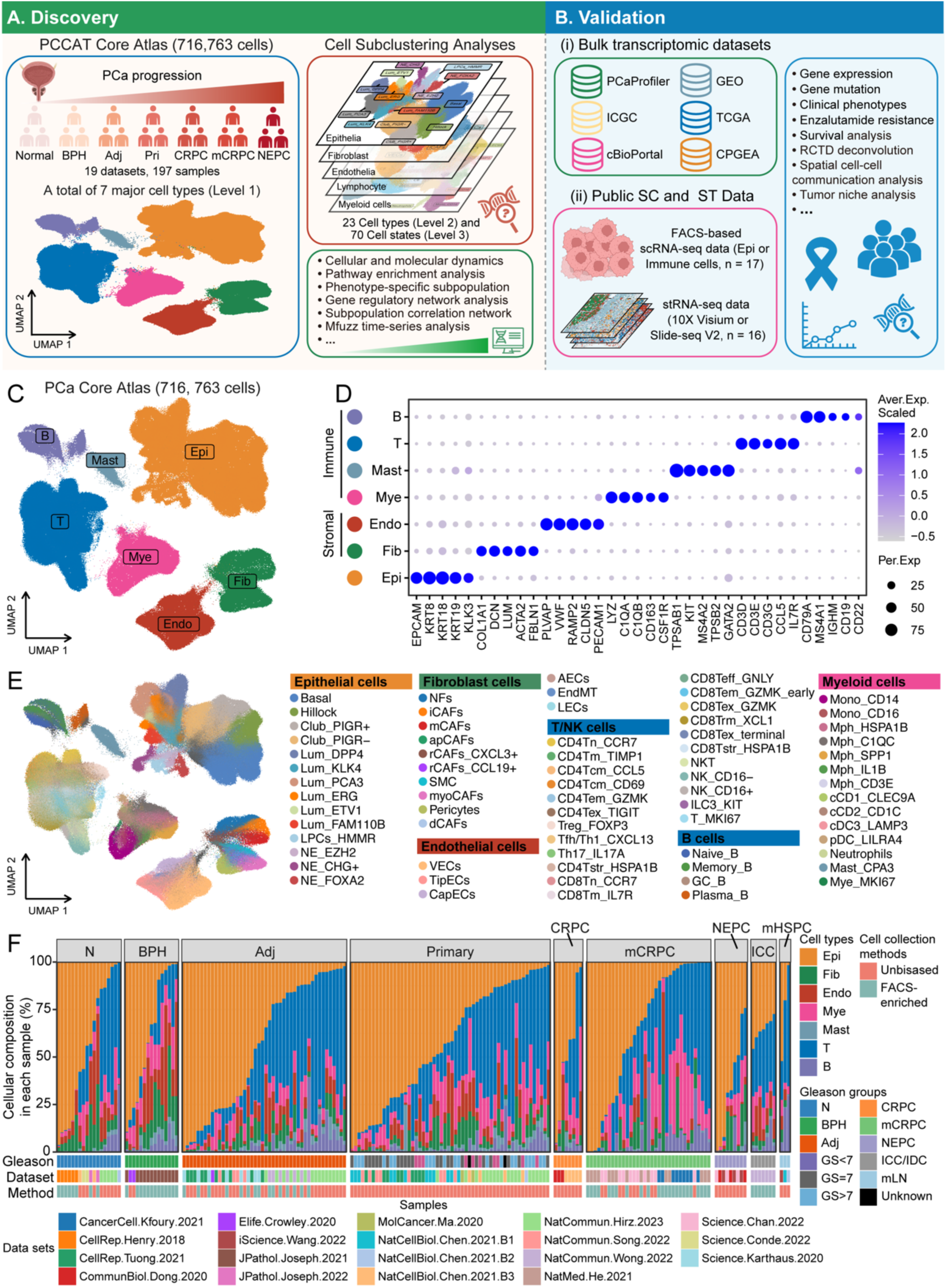
PCCAT profile of 716,763 cells from 197 human samples covering various tumor stages of PC. A and B. Analysis workflow of this study. C. UMAP plot of the 716,763 single cells in PC core atlas, colored by 7 major cell types. Epi, epithelial cells; Fib, fibroblasts; Endo, endothelial cells; Mye, myeloid cells. D. Dot plot of representative marker genes for each major cell type. Bubble size indicates the percentage of cells expressing a specific gene in a given cell cluster (Per.Exp), and color depicts scaled average expression (Aver.Exp.Scaled). E. UMAP plot displaying the 70 cell clusters in the PC core atlas. F. Composition of each cell type in each sample, grouped by disease progression. N, normal; BPH, benign prostate hyperplasia; Adj, normal tissues adjacent to the tumors; GS, Gleason score.

By benchmarking 11 top-performing batch-correction and integration tools (Figures S3 and S4), we determined that the BBKNN algorithm was the most suitable for generating the PCCAT and was thus chosen for further analyses. Overall, we constructed a core atlas integrating 716,763 high quality single cells, which are annotated to 7 major cell types, 23 refined cell subtypes, and 70 transitional cell states based on previously established canonical single-cell markers^39^ (Figures 1C-1E). The 7 major cell types exhibit wide ranges in relative abundance across 197 samples representing diversity within and across disease stage groups (Figure 1F).

### Characterizing the heterogeneity of epithelial cells during PC progression

We identified six distinct epithelial cell subtypes at annotation level 2, encompassing further differentiation into 14 cell states at level 3 annotation (Figures 2A, and S5A-S5C). Each cell received a malignancy score based on assessing carcinoma vs benign cell genotypes using inferCNV^40^ to determine structural genomic alterations associated with neoplasia (Figures S5D-S5F). Consistent with previous reports^18, 35, 41^, the basal, hillock and club cell types are identified as nonmalignant, predominantly originating from non-tumor samples (Figures 2B-2D). Cells annotated as luminal, lineage plasticity-like cells (LPCs) and NE cells were primarily associated with malignant cells derived from tumor samples (Figures 2B-2D). Moreover, as PC progressed, we noted an elevated malignancy classification rate among the epithelial cell populations, coupled with a reduction in the expression of immune checkpoint genes and transcripts encoding Major Histocompatibility Complex (MHC) class I proteins (Figures 2D, S5G, and S5H), suggesting that the downregulation of MHC class I expression is associated with increased invasiveness and progressively higher carcinoma grades. The expression levels of PD-1 and CTLA-4 ligand genes, namely *CD274* (PD-L1) and *CD80/86* (B7-1/2), remained relatively low across all stages of PC, which is consistent with the limited responsiveness to PD-1 or CTLA-4 blockade reported in clinical studies^42, 43^.

**Figure 2.**
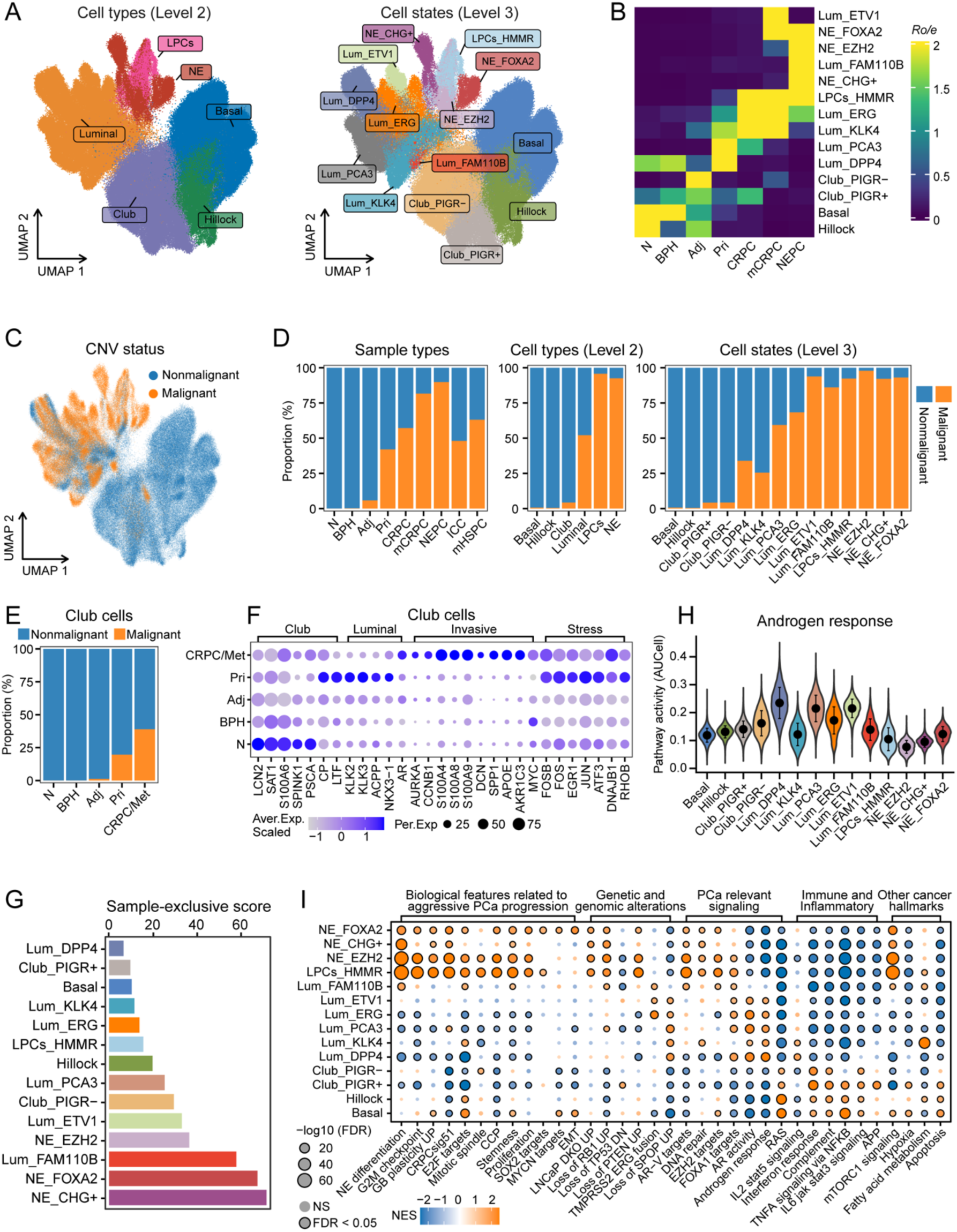
Characteristics of epithelial cell subpopulations during PC progression. A. UMAP plots showing all epithelial single cells, colored by cell types (left) and cell states (right). LPCs, lineage plasticity-like cells; NE, neuroendocrine cells. B. Tissue preference of each epithelial cluster estimated by Ro/e score. C. UMAP plots showing the distribution of malignant cells inferred by using inferCNVpy. D. Bar plots showing the proportion of malignant cells for each sample subtype (left), cell type (middle), and cell states (right) in epithelial cells of PC. E. Bar plots showing the proportion of malignant cells for each sample subtype in Club cells of PC. F. Dot plot showing the dynamic expression of representative genes associated with indicated epithelial features in Club cells during PC progression. G. Individual sample occupancy (sample-exclusive score) for each epithelial cell cluster. A high sample-exclusive score for a cell cluster indicates a low degree of sharing across different samples. H. AUCell enrichment analysis comparing the activities of androgen response among different epithelial cell clusters. I. Gene set enrichment analysis (GSEA) of transcriptome dynamics in different epithelial cell subclusters. NES, normalized enrichment score; NS, no significant.

As predominant nonmalignant epithelial cell types, basal and hillock cells showed significantly higher frequencies in normal and BPH prostate tissues, while club cells exhibited a preference for tumor adjacent tissues (Figures S5I-S5K). Moreover, we identified similarities between basal and hillock cells, as well as between club and luminal cells (Figures S6A and S6B), indicating that hillock and club cells may serve as basal and luminal cell progenitors, respectivley^30, 35^. Furthermore, these nonmalignant epithelial cell types exhibit PC-enriched luminal-like cell states characterized by upregulated AR signaling in primary PC (Figure S6C). This phenomenon is not limited to basal and club cells^30^ but also extends to hillock cells, suggesting a widespread loss of original cell identity and acquisition of luminal-like states among nonmalignant cell types in early PC tissues. Notably, club cells exhibited an elevated malignancy classification rate by inferCNV analysis with PC progression, and increased expression of several well-known invasive and stress-related genes at advanced stages, such as *AURKA*, *CCNB1*, *S100A4*, *FOS,* and *DNAJB1* (Figures 2E, 2F, and S6D). This suggests that club cells may not only be associated with early phases of carcinogenesis^30, 44, 45^ but also contribute to tumor progression.

Among epithelial cells involved in malignant progression, Lum_DPP4 and Lum_KLK4 exhibited the lowest individual sample-exclusive score^46^, representing prevalent cell states in both normal prostate and tumor tissues (Figures 2G). The two subpopulations also demonstrated a low malignancy classification rate (Figure 2D) and were characterized by distinctly opposing androgen response features (Figure 2H). Notably, Lum_DPP4 cells highly expressed classical AR target genes (e.g., *DPP4*, *MSMB* and *ACPP*), displayed heightened features associated with luminal-AB cells^44^, and AR activity, and were significantly reduced in frequency in the CRPC phase (Figures 2I and S7A-S7D). This suggests that they might represent castration-sensitive low-malignancy luminal cells. On the other hand, Lum_KLK4 cells, previously identified as a ubiquitous population situated between KLK5+ basal cells and KLK3+ luminal cells^23^, displayed elevated *KLK4* expression and low expression of *AR* and its associated target genes (Figure S7A). Concurrently, Lum_KLK4 cells were significantly enriched during PC progression (Figures 2B and S7B), and showcased attenuated androgen responsiveness and AR activity (Figures 2H and S7D), potentially representing a subset of a castration-resistant luminal cells.

Epithelial cells annotated as Lum_PCA3 and Lum_ERG represented the predominant malignant luminal cells in both localized PC and CRPC, characterized by activated PC relevant signaling, such as Luminal-AB features, androgen responsiveness, and FOXA1 signaling (Figures 2B,2H, and S7A-S7D). Moreover, Lum_ERG cells were characterized by high *ERG* expression, along with the ERG+ tumor markers *PCA3* and *AMACR*, and activated TMPRSS2-ERG fusion pathway^30^ (Figures 2I and S7A), suggesting the presence of ERG fusion events. In addition, we identified four NEPC-related malignant cell states each associated with poor prognosis (Figures 2A, 2B, and S7A-S7C). First, LPC_HMMR cells comprised a rare population of AR+ HMMR+ CHGA-malignant luminal cells pre-existing in primary tumors and harboring luminal-neuroendocrine trans-differentiation^41^. These cells are distinguished by their low AR signaling and elevated *HMMR* and *EZH2* expression (Figures 2H and S7A) and enhanced lineage plasticity activity (Figures 2I and S7C-S7E), such as expression of EZH2 targets, TP53/RB1-inactivated UP signature, and CRPCsig51 (a gene signature related to a pre-existing castration resistant cell type) ^41, 47, 48, 49, 50^. Furthermore, this cell subpopulation exhibited relatively high expression of genes related to epigenetic modifiers^11^, such as *DNMT3A*, *EZH2* and *SUZ12*, and others related to stemness and NE progression^39, 41, 51, 52, 53, 54^, such as *EZH2*, *E2F1*, *AURKA*, *XPO1*, *SOX2*, and *NKX2-1* (Figure S7E).

Interestingly, we observed a lower sample-exclusive score for LPCs_HMMR compared to other three NE subclusters, suggesting this cell population is more widely present in samples with PC (Figure 2G). We also identified three classic NE-like subpopulation: NE_EZH2, NE_CHG+ and NE_FOXA2 that expressed classic NE markers and were enriched in advanced PC (Figures 2B and S7A). Remarkably, a distinctive feature of NE_FOXA2 cells involves the up-regulation of EMT-related genes, such as *COL1A2*, *FGF3/19/20*, *VIM* and *TWIST1* (Figure S7E), supporting a potential mechanism for the high rate of metastasis in NE-like tumors. Taken together, these results depict a dynamic heterogeneous landscape of both non-malignant and malignant epithelial cells during PC progression (summarized in Table 1).

**Table 1.**
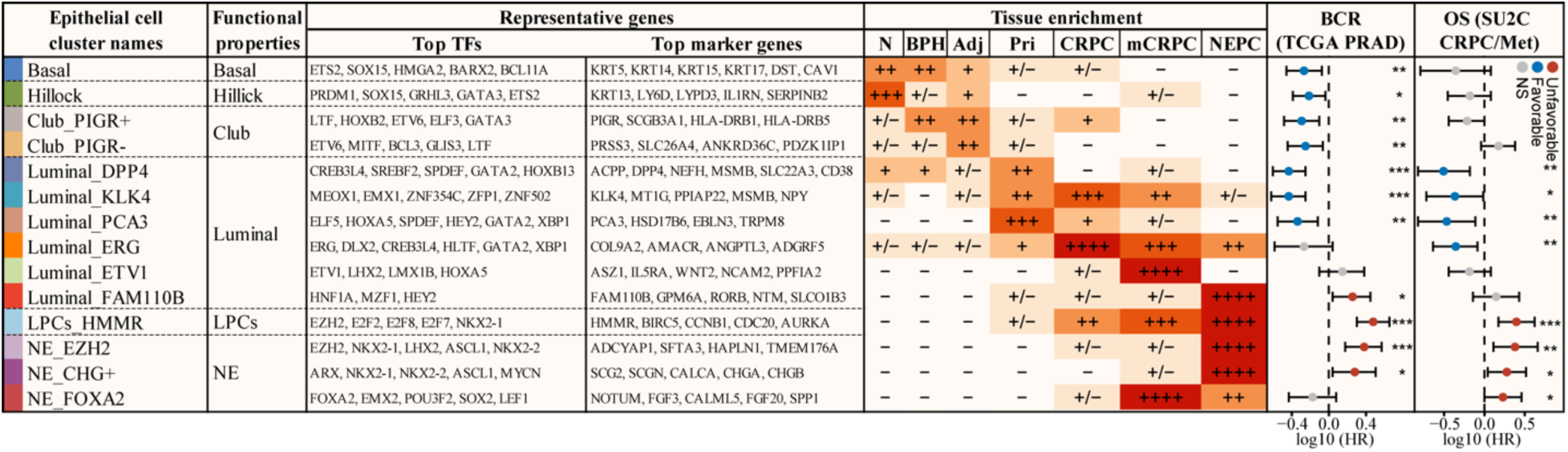
Summary of epithelial cell subpopulation during PC progression. Comprehensive overview of the characteristics of epithelial subsets derived from prostate cancer patients, encompassing functional properties, representative genes, tissue enrichment and clinical relevance. ++++, Ro/e > 5; +++, 2 < Ro/e ≤ 5; ++, 1.5 < Ro/e ≤ 2; +, 1 < Ro/e ≤ 1.5; +/−, 0.1 < Ro/e ≤ 1; −, 0 ≤ Ro/e ≤ 0.1. *, P < 0.05; **, P < 0.01; ***, P < 0.001; NS, no significant. N, normal; BPH, benign prostate hyperplasia; Adj, normal tissues adjacent to the tumors.

### Integration of bulk RNA-seq data reveals genotype-phenotype associations within epithelial cell subpopulations

The scRNA-seq method provides quantitative insights into the cellular diversity within the TME. However, most scRNA-seq studies evaluating PC are limited by the absence of both cancer genotype information and survival outcome data. In contrast, substantial bulk RNA-seq datasets like TCGA PRAD^7^, SU2C CRPC/Met^55^, and PCaProfiler^11^ are accompanied by rich orthogonal information that includes genomic mutation status, biochemical recurrence (BCR), overall survival (OS), treatment responses and histological assessments such as Gleason grade. To leverage these data types, we employed the SuperCell algorithm to aggregate ∼267 k individual epithelial cells into ∼64 k high-resolution epithelial metacells separately within each sample (Figure S8A), which mitigates technical differences within single cells and enhances cell quality^56^. Building on this epithelial metacell atlas, we then applied the Scissor computational method^57^, to evaluate the association between atlas-derived cell-types and clinical phenotypes from the bulk RNA-seq datasets.

The high-resolution of the epithelial metacell PC atlas enabled an in-depth analysis of epithelial subpopulations that associate with tumorigenesis, tumor progression, patient prognosis, enzalutamide resistance, and specific somatic genomic alterations such as *TMPRSS2-ERG* fusion, mutations in *SPOP*, *PTEN*, *FOXA1*, *TP53* or *RB1* (Figures 3A, 3B, and S8-S11). The pathway enrichment and spatial analyses further confirmed the accuracy of the key phenotype-related metacells and molecular features identified by the Scissor algorithm (Figures 3C and S12). For example, we identified a correlation between the Lum_ERG subpopulation and *TMPRSS2-ERG* fusion (Figures 3A-3C). Loss of *RB1* or *TP53*, representing established drivers of lineage plasticity and therapeutic resistance^49, 51, 58^, were linked to cell types annotated as NE_EZH2, NE_CHG+, NE_FOXA2 and LPCs_HMMR (Figures 3B, S9C, and S9D). Moreover, we found that Lum_DPP4, Lum_PCA3, and Lum_ERG subpopulations were associated with favorable prognosis in both early and advanced PC (Figures 3B and S10A), whereas Lum_ETV1, Lum_FAM110B, LPCs_HMMR and the three NE subpopulations were correlated with tumor progression and unfavorable prognosis (Figures 3B, S9E, and S10A). Based on these findings, we further combined BCR-related and OS-related metacells to identify the key cell states and molecular characteristics associated with PC prognosis (Figures S10B-S10E). Notably, the results suggested that metacells linked to poor prognosis exhibited enhanced stemness, mTORC1 signaling, lineage plasticity, and cell cycle related pathways, while showing reduced expression of MHC related genes (Figures 3E and S10C-S10E). Subsequently, we developed a signature that associated with BCR and OS (Table S2), which annotated the spatial distribution of high-risk versus low-risk tumor niches and exhibited robust outcome prediction (Figures 3D-3G, S10F, and S10G).

**Figure 3.**
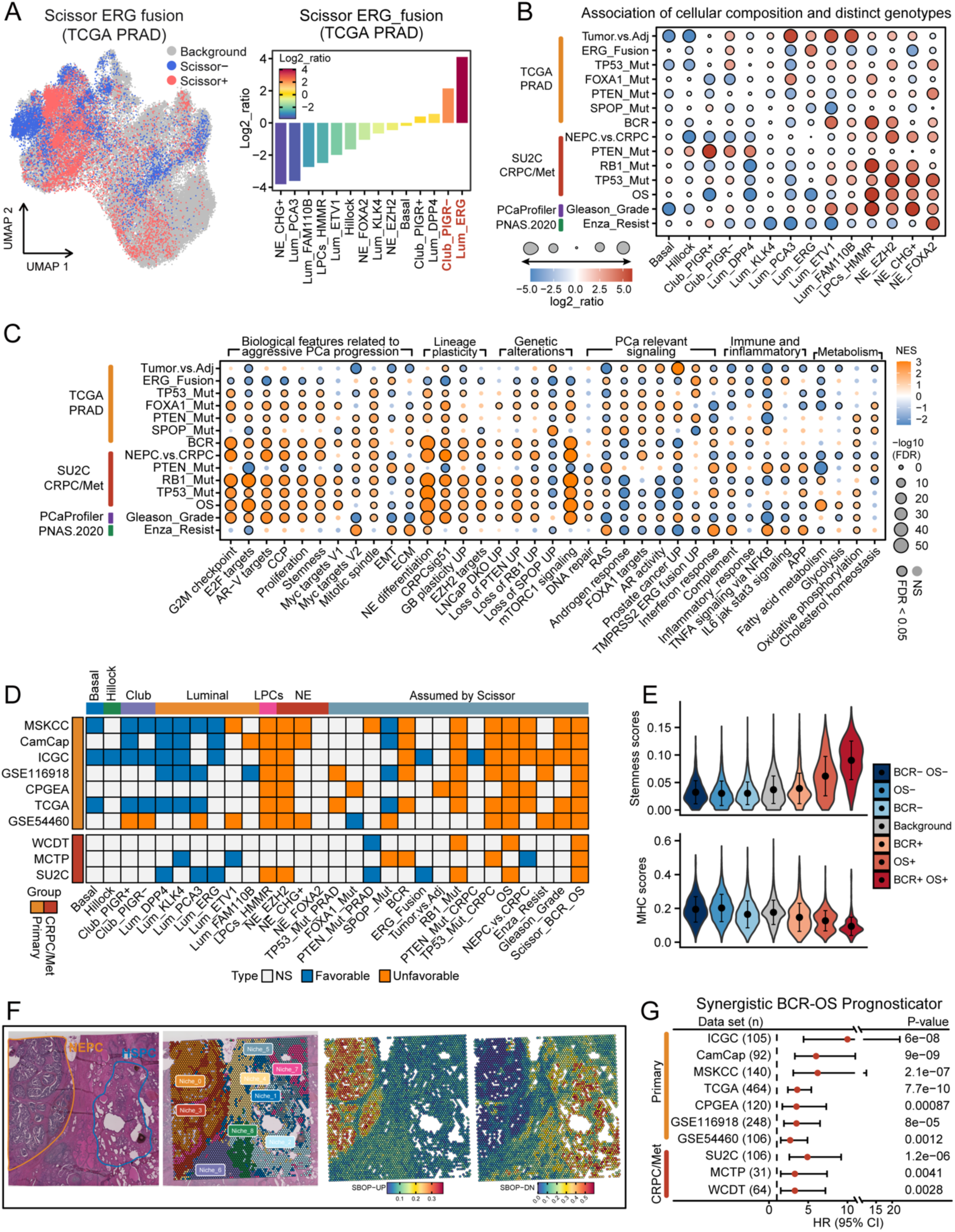
Association between epithelial cellular composition and distinct genotypes, prognosis as well as treatment resistance. A. Scissor analysis of ERG fusion from TCGA PRAD bulk RNA-seq data with epithelial cells. UMAP plots indicate the position of cells positively (red) or negatively (blue) associated with ERG fusion. A log2 ratio >0 indicates a positive association with ERG fusion, respectively. B. Dot plot showing log2 ratio between the indicated phenotype’s Scissor positive epithelial cells and Scissor negative epithelial cells for each epithelial cluster. A log2 ratio >0 indicates a positive association with mutation, better survival or treatment resistance, respectively. C. GSEA analysis of transcriptome dynamics in Scissor positive epithelial cells and Scissor negative epithelial cells related to indicated phenotypes. NES, normalized enrichment score; NS, no significant. D. Univariate Cox regression analysis for each epithelial cluster and the Scissor-derived signatures related to the indicated phenotype. Survival index using biochemical recurrence (BCR) for primary PC and overall survival (OS) for CRPC/Met PC. E. The distribution of pathway activities for stemness (upper) and MHC activity (lower) among epithelial cells related to different prognostic subpopulations. F. The spatial feature plot showing the score distribution of Scissor-derived BCR and OS associated prognosticator (SBOP). G. Forest plot showing survival analysis of SBOP signature for BCR in primary PC cohorts or OS in CRPC/Met cohorts. HR, Hazard ratio; 95% CI, 95% confidence interval.

Next, we conducted a detailed exploration of the molecular characteristics associated with response to enzalutamide treatment. Our findings revealed an association between the NE_FOXA2 subpopulation and enzalutamide resistance, while the majority of luminal subpopulations associated with responsiveness to enzalutamide treatment (Figure S11A). Moreover, pathway enrichment analysis showed that metacells associated with enzalutamide resistance exhibited significantly activated interferon response, Myc targets, extracellular matrix (ECM), and epithelial-mesenchymal transition (EMT) signaling, alongside suppressed androgen response and FOXA1 target signaling (Figures S11B and S11C). Interestingly, enzalutamide resistance-related meta cells had a high activity of MHC signaling (Figure S11C). These results indicate a correlation between low AR activity, high MHC expression and a stemness epithelial program with enzalutamide resistance, further reinforcing our previous findings from bulk transcriptome data^3^. Building on this perspective, we further developed an enzalutamide treatment resistance gene signature (Table S2), demonstrating strong predictive capability for poor response in both human PC samples and several drug-resistant PC cell lines (Figures S11D-S11J).

### Stromal cell subtypes associate with malignant features of advanced PC

There is increasing evidence that stromal cells contribute to the development of PC recurrence, metastasis and therapy resistance^59, 60, 61^. Here, we identified 4 major stromal cell lineages, comprising endothelial cells, fibroblasts, myofibroblasts, and pericytes, along with 16 clusters representing sublineages with distinct functional roles (Figures 4A, 4B, and S13A-S13C). Scissor analyses further unveiled distinct associations of tumor mutations (including *PTEN*, *TP53*, and *SPOP*) and *ERG* fusion with stromal components (Figure S13D). Interestingly, we observed an association between endothelial cells and Gleason grade (Figure 4C), suggesting a connection to disease progression in PC. Consistently, Gene Ontology (GO) analysis revealed that up-regulated differentially expressed genes (DEGs) in genotype-associated endothelial cells were enriched for several pathways associated with malignant progression, involving endothelial cell differentiation, migration, and angiogenesis (Figure S13E). Moreover, survival analyses confirmed that the signature genes of endothelial cells and pericytes were both correlated with BCR in multicenter PC cohorts (Figure 4D).

**Figure 4.**
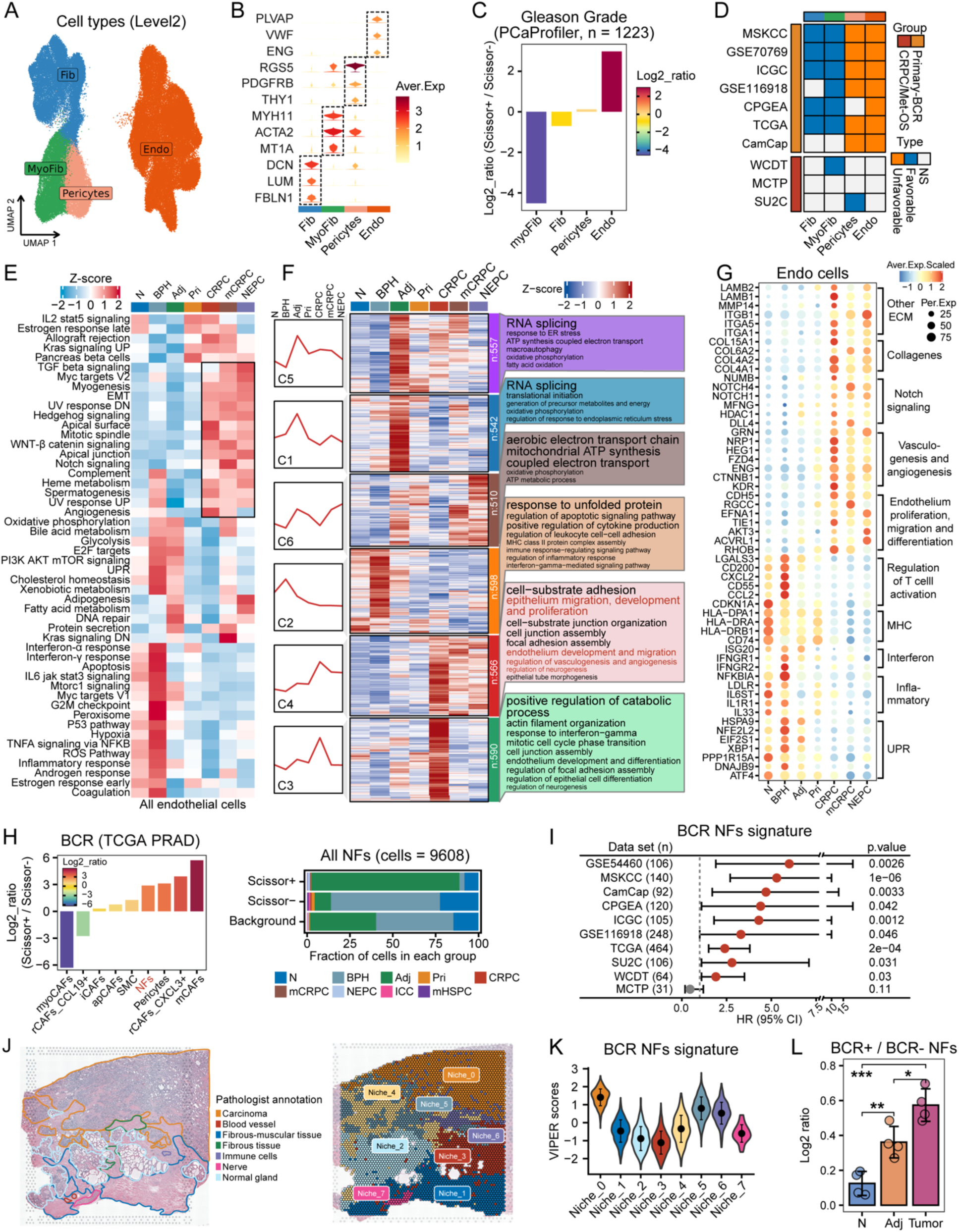
Characteristics of stromal cells during PC progression. A. UMAP plot of all stromal cells, colored by cell types. Fib, fibroblasts; MyoFib, myofibroblasts; Endo, endothelial cells. B. Violin plot showing representative marker genes for each stromal cell type. C. Identification of Gleason grade related metacells using Scissor linear regression analysis with PCaProfiler bulk dataset. Log2 ratio >0 indicates a positive association with high PC grade. D. Univariate Cox regression analysis for each stromal cluster. Survival index using BCR for primary PC and OS for CRPC/Met PC. E. Heatmap showing the activity remodeling of functional hallmark pathways for endothelial cells as disease progression. F. Mfuzz time-series analysis illustrating the patterns of dynamic transcriptional changes in endothelial cells during PC progression. The line graph and heatmap depict the change patterns of six gene clusters as disease progression. And the word cloud displays enriched GO terms for corresponding gene cluster. G. Dot plot showing the dynamic expression of representative genes associated with indicated pathways during disease progression. H. Bar plot showing BCR related fibroblast metacells using Scissor Cox analysis with TCGA PRAD bulk dataset (left), and proportions of each normal fibroblast metacells in BCR related Scissor groups (right). Log2 ratio >0 or Scissor+ indicate a positive association with poor prognosis. Scissor-indicates a positive association with good prognosis. I. Forest plot illustrating survival analysis of the BCR^+^ NFs signature derived from DEGs between Scissor+ and Scissor-metacells associated with BCR. HR, Hazard ratio; Error bars denote 95% confidence interval (CI). J. The pathological annotation (left) and cell niches (right) of stRNA-seq data from the invasion leading edge of primary PC. K. The spatial distribution of the BCR^+^ NFs signature scores in the invasion leading edge section. L. Bar plot showing the distribution of the BCR^+^ NFs among different PC groups, based on all fibroblast cells. N, normal; Adj, normal tissues adjacent to the tumors. Kruskal-Wallis test with Bonferroni correction. *, p < 0.05; **, p < 0.01; ***, p < 0.001. Error bars denote SD (K and L).

To further investigate the molecular remodeling of endothelial cells by cancer cells^18^, we first performed AUCell enrichment^62^ using 50 Hallmark gene sets to analyze the alterations in pathways of endothelial cells among different PC stages. Intriguingly, we found that several immune-related pathways, such as inflammatory response, IL6/JAK/STAT3 signaling, and interferon response, were enriched in endothelial cells from BPH samples, while a subset module of hallmark pathways involving crucial tumor endothelial cell signals were enriched in advanced PC, such as TGF-β signaling, Myc targets V2, WNT-β catenin signaling, Notch signaling, and angiogenesis (Figure 4E). In addition, Mfuzz time-series analysis^63^ identified six gene clusters (GC1-6) exhibiting distinct molecular dynamics and functions in endothelial cells during PC progression. For example, GC2 exhibited high expression levels in non-tumor samples and decreased in a stepwise fashion with PC progression. This GC2 cluster was enriched for genes involved in the unfolded protein response (UPR) and several immune-related pathways, such as positive regulation of cytokine production, MHC class II protein complex assembly, and regulation of inflammatory response (Figures 4F, 4G, and S13F). Moreover, the expression levels of genes in GC4 were upregulated in advanced PC, with enriched functions related to the ECM, endothelial cell development, migration, and angiogenesis, as well as epithelial cell migration, proliferation, and neurogenesis (Figures 4F, 4G, and S13F), highlighting their connection to malignant progression. Together, these results support the existence of an interactive relationship between cancer cells and the induction of distinct endothelial cell states that may facilitate disease progression.

We next sought to further characterize the 105,049 high quality single cells that comprised the four major stromal cell types identified across all of the atlas samples. We performed a re-cluster analysis and partitioned the four major stromal types into 16 subtypes, including normal fibroblasts (NFs), 7 cancer associated fibroblasts (CAFs), smooth muscle cells (SMCs), pericytes, and 6 endothelial cell subpopulations (Figures S13G-S13I). These stromal cell subtypes exhibited heterogeneous tissue preference and molecular features during PC progression (Figures S13J and S14A-S14D). For example, matrix CAFs (mCAFs), pericytes, and tip-like endothelial cells (TipECs) were all enriched during PC progression and accounted for the majority of the stromal populations (Figures S13J and S14A). Among them, mCAFs were characterized by low levels of *ACTA2* (*α-SMA*) but high levels of ECM signatures, including collagen molecules (*COL1A1*, *COL3A1*, and *COL5A1*), and PC bone metastasis related markers^61^ (*POSTN*, *ASPN*, and *FSCN1*) (Figures S13H and S14C). Notably, enrichment analyses showed that this subcluster was involved in several pathways potentially contributing to disease progression, such as ECM, regulation of angiogenesis, bone development, and EMT signaling (Figures S14B and S14D). Furthermore, we observed a prominent correlation between the mCAFs and tumorigenesis, high Gleason grade, as well as poor prognosis (Figures S14E, S14F, and 4H). As for pericytes, we found that this subpopulation mainly expressed pro-angiogenic factors such as *RGS5*, *NOTCH3*, and *CD146* (*MCAM*), as well as PC bone metastasis related markers^61^ such as *PDGFRB* and *PMEPA1* (Figures S13H and S14C). GO analysis of pericytes indicated significant enrichment for genes involved in endothelial cell migration and vascular transport, consistent with their pro-angiogenic features (Figure S14B). Lastly, TipECs, primarily found in tumor tissues across various cancer types^64^, mainly expressed tip cell related markers such as *PLVAP*, *INSR*, and *ESM1*, and showed significant enrichment for endothelial cell migration, angiogenesis, and oxidative phosphorylation (Figures S13I and S14B). Collectively, these findings underscore adverse prognostic cell states and distinctive features of the stromal components present in advanced PC, including ECM remodeling, heightened EMT, and active angiogenesis signaling, when compared to localized PC and non-tumor tissues.

### Identification of a minority population of normal fibroblasts associated with BCR (BCR^+^ NFs) from tumor-adjacent tissues

Compared to CAFs, which constitute the most abundant component in the PC TME^61^, we found that cells classified as SMCs and NFs were enriched in non-tumor tissues (Figures S13J and S14A). Notably, SMCs expressed higher levels of typical smooth muscle-related markers, such as *α-SMA*, *MYH11* and *MUSTN1*, and expressed genes involved in muscle cell differentiation (Figures S13H and S14B). NFs exhibited a combination of peri-epithelial fibroblast characteristics (*PTGDS* and *PTGS2*) and interstitial fibroblast features (*FGF2*, *CCK*, and *C7*) (Figure S13H), two novel fibroblast subtypes recently identified in the human prostate^29^. GO enrichment analyses showed that this fibroblast cell subcluster expressed genes involved in the UPR, inflammatory response, and leukocyte recruitment process (Figures S14B and S14D).

Consistent with the presumed anti-tumorigenic effects for NFs^61^, our findings suggested a positive correlation between NFs and a favorable prognosis in patients with primary PC (Figures S14F). However, employing Scissor analysis, we noted a positive association between a small population of NFs and BCR (hereafter BCR^+^) (Figure 4H). Further analysis revealed that the BCR^+^ NFs, which were associated with unfavorable survival, were mainly derived from tumor-adjacent tissue samples, accounting for over 80% of the total BCR+ NFs (Figure 4H). Recent studies have demonstrated that the tumor-adjacent tissue represents a unique intermediate state between healthy tissue and tumor, reflecting a field-effect capable of promoting further neoplasia, and also potentially harboring oncogenic events^65, 66^. In PC, alterations in the stromal gene signature may precede those in the epithelial gene signature during PC initiation and progression^59^.

To gain deeper insight into rare BCR^+^ NF subtype, we first performed differential expression analysis between BCR^+^ NFs and BCR^—^ NFs, leading to the identification of a BCR^+^ NF gene signature (Table S2). Survival analysis further demonstrated a stable association between BCR^+^ NF and poor prognosis in multiple multicenter cohorts (Figures 4I). In addition, GO enrichment analyses revealed that up-regulated DEGs in BCR^+^ NFs were enriched for several energy conversion and biosynthesis related pathways, such as cellular respiration, oxidative phosphorylation, and ATP metabolic process, as well as exosomal secretion (Figure S15A). Up-regulated DEGs in BCR^—^ NFs were also enriched for several UPR and immune regulation related pathways, such as response to unfolded protein, antigen processing and presentation, leukocyte cell-cell adhesion, and positive regulation of T cell activation (Figure S15A). Lastly, SCENIC analysis^62^ identified transcription factors (TFs) that exhibited high regulon specificity in BCR^+^ NFs and were associated with malignant tumor progression^67, 68, 69^, including HOXB2, CREB3, CREB5, and HIF1A. Meanwhile, BCR^—^ NFs exhibited elevated expression of several stress and immune related TFs, such as ATF6, FOSB, IRF9, and CEBPD (Figure S15B).

As additional validation, we investigated the spatial distribution and abundance of BCR^+^ NFs by integrating data from multiple stRNA-seq PC tissue slices with the fibroblast single-cell atlas generated here. First, using a primary PC tissue section annotated as ‘invasion leading edge’, both signature enrichment analysis and the RCDT deconvolution algorithm revealed that BCR^+^ NFs were mainly found in the tumor zone (Niche_0) and the tumor invasive zone (Niche_5 and Niche_6) (Figures 4J, 4K and S15C). In contrast, BCR^—^ NFs were predominantly enriched in the non-tumor zone (Niche_1-4, and Niche_7) (Figure S15C). Additionally, we found a stepwise increase in the enrichment of the gene signature of BCR^+^ NFs as PC progressed from normal to leading-edge to tumor regions (Figure S15D). Similar results were further validated in nine Slide-seqV2 stRNA-seq slices (Figures 4L, S15E, and S15F). Taken together, these results reveal that a minority population of BCR^+^ NFs may pre-exist in PC adjacent tissues, potentially influencing tumor dissemination and recurrence.

### Characteristics of lymphocytes in the prostate TME

In addition to resident stromal cells, tumor-infiltrating lymphocytes are key elements of the prostate TME and have the potential to influence adverse PC behaviors including invasion, metastasis and treatment responses^70, 71, 72^. Of the 716,763 cells analyzed, a total of 30,860 (4.3%) classified as B cells comprised of two major types, B lymphocytes and plasma B cells, with a further subdivision into four B cell states, namely naïve B, memory B, germinal center (GC) B, and plasma B, according to canonical markers (Figures 1D, S16A and S16B). Naïve B, memory B, and GC B showed high expression of MHC class II genes, while plasma B highly expressed *SDC1* (CD138), *TNFRSF17* (BCMA), *MZB1*, *JCHAIN* with no/low expression of MHC class II genes (Figure S16C). Tissue preference analysis further showed that plasma B cells had a strong distribution preference in tumors relative to benign tissue samples, while GC B cells were mainly enriched in mCRPC and NEPC (Figure S16D).

A total of 204,246 cells were classified as T/Natural killer (NK) cells, which were subclassified into five major cell types (Figures 5A, S17A, and S17B): CD4 T, CD8 T, cycling T, NK/NKT, and Type 3 innate lymphoid cells (ILC3s). These subclassification were further grouped into 18 T cell states and four innate lymphoid cell states according to established markers and curated gene signatures^70, 71, 72^ (Figures 5A, 5B, and S17C; Table S3). The cycling T cells showing higher proliferation activity were also designated as T_MKI67, and their proportions significantly expanded during the progression of PC (Figures 5C and S17B). Interestingly, cycling T cells exhibited elevated expression of exhaustion-related regulators^70^ and expressed a slightly lower exhaustion signal compared with the terminally-exhausted CD8+ T cells, but exhibited low to absent expression of *TNF*, *TXB21* and *IL2* (Figure S17D). This implied that cycling T cells may lack sufficient cytotoxic and pro-inflammatory capacity^72^, potentially contributing to the attenuation or suppression of immune responses.

**Figure 5.**
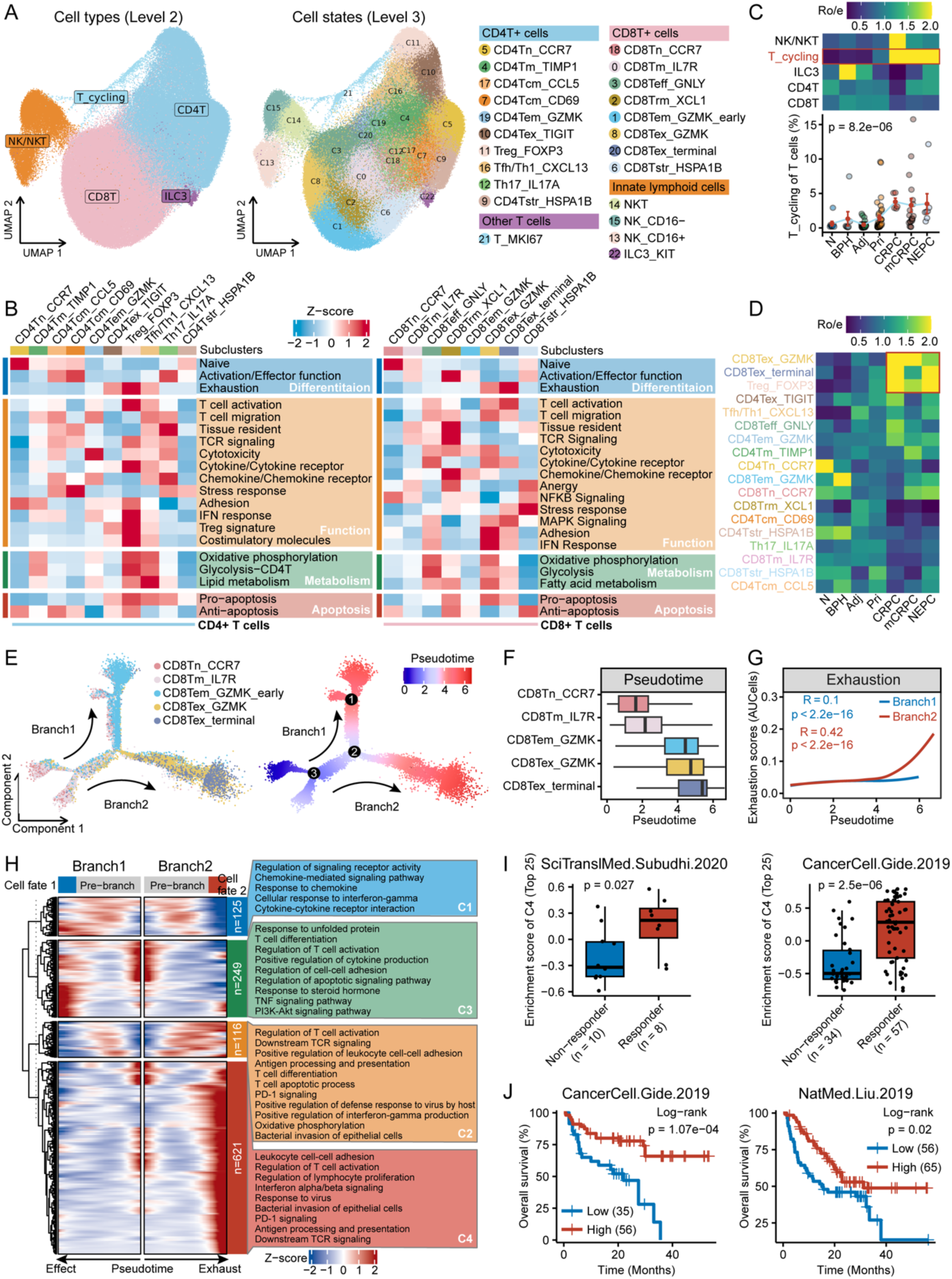
Characteristics of T and NK cells during PC progression. A. UMAP plots of all T and NK cells. B. Heatmap illustrating expression of curated gene signatures across CD4+ T (left) and CD8+ T (right) cell clusters. C. Tissue preference of each T/NK cell type estimated by Ro/e score (upper), and scatter diagram showing the frequency change of T_cycling cells with PC progression (lower). D. Tissue preference of each T cell cluster estimated by Ro/e score. E. Monocle 2 trajectory analysis of CD8+ T cell differentiation revealing two main divergent trajectories. Cells are color coded for their corresponding cell clusters (left) or pseudotime (right). F. Boxplot showing the diffusion pseudotime distribution of CD8+ T cells in the two major paths from naïve T cells to exhaustion. G. Two-dimensional plots showing expression scores for exhaustion gene signatures in cells of path branch 1 (blue) and path branch 2 (red) respectively, along the inferred pseudotime. H. The DEGs (rows) along the pseudotime (columns) are hierarchically clustered into four gene clusters. The representative annotated pathways of each gene cluster are provided. I. Enrichment scores for the C4 gene cluster signature, composed of the top 25 genes, in individual patients from an immunotherapy dataset of mCRPC (left) and a published immunotherapy dataset of metastatic melanoma (right). J. Kaplan–Meier survival analysis illustrating overall survival outcomes for patient groups with low and high scores of the C4 gene cluster signature, comprising the top 25 genes, in two immunotherapy cohorts of metastatic melanoma.

We further characterized the molecular features, tissue preference, and dynamic changes in cell frequency of each T cell subpopulation during PC progression (Figures 5B, 5D, S17E, and S17F). Notably, three immunosuppression-related T cell clusters including Treg_FOXP3, CD8Tex_GZMK, and CD8Tex_terminal cells, were significantly enriched with PC progression (Figures 5D and S17E), suggesting that a suppressive TME exists in advanced PC. Lastly, SCENIC analysis uncovered several established^70, 71, 72, 73^ and novel TFs in Treg_FOXP3 (e.g., ETV7, SOX, and GATA3), CD8Tex_GZMK (e.g., IRF9, EOMES, and STAT1), and CD8Tex_terminal (e.g., ETV1, AHR, and GATA3) cells (Figure S17F).

The comprehensive transcriptomic data derived from a large number of T cells provides insights into the functional states and the potential inter-relationships between these cells. First, we applied the Monocle 2 algorithm to order CD8+ T cells in pseudotime to assess their developmental trajectories (Figure 5E). The results revealed the aggregation of five CD8+ T clusters based on gene expression similarities with a pseudotime projection starting with CD8Tn_CCR7 (naïve CD8+ T cells), followed by CD8Tm_IL7R (memory CD8+ T cells). The trajectory then diverged into branches representing distinct cell differentiation fates: one leading to CD8Tem_GZMK (effector memory CD8+ T cells), while the others led to CD8Tex_GZMK (exhausted CD8+ T cells) and CD8Tex_terminal (terminal exhausted CD8+ T cells) (Figures 5E-5G). Along the trajectory, we identified four gene clusters (Table S4), where C1 and C3 were associated with branch 1 (naïve to effector memory), while C2 and C4 were associated with branch 2 (naïve to terminal exhaustion) (Figure 5H). Notably, C2 exhibited a continuous expression distribution between CD8Tn_CCR7 and CD8Tex_terminal cells (Figure 5H), indicating a pre-exhausted CD8+ T cell transition. Furthermore, enrichment analysis revealed involvement in several pro-exhaustion-related pathways^73^, encompassing the regulation of T cell activation, downstream T cell receptor (TCR) signaling, PD-1 signaling, as well as factors related to persistent antigen exposure, including antigen processing and presentation, virus infection, and bacterial invasion (Figure 5H). The genes in C4 were predominantly expressed in CD8Tex_GZMK and CD8Tex_terminal cells and enriched for terminal exhaustion-related pathways^72, 73^, such as regulation of lymphocyte proliferation and interferon related signaling (Figure 5H).

To further refine the characteristics of T cells expressing the C2 and C4 gene clusters, we identified the top 25 genes positively correlated with pseudotime in branch 2 that distinguished C2 and C4 (Figures S18A). The C2’s top genes were predominantly associated with cytotoxicity (e.g., *GZMA*, *GZMH*, and *NKG7*), MHC (e.g., *HLA-DPB1*, *HLA-DRB5*, and *HLA-C*), and T cell activation signaling (e.g., *CD2* and *CD27*), while the C4’s top genes encode well-known exhaustion markers^70, 74^ (e.g., *CXCL13*, *CTLA4*, *DUSP4*, *HAVCR2*, *TOX*, and *RBPJ*). A strong positive correlation between the two gene signatures and exhaustion gene signature scores^74^ was also observed using a large bulk RNA-seq PC cohort (Figure S18B). Notably, the gene signature of C4 exhibited robust predictive capabilities for assessing responses to immune checkpoint blockade (Figures 5I and S18C) and survival outcomes (Figures 5J and S18D) across six independent immunotherapy cohorts (Table S1). Collectively, these findings characterize diverse lymphocyte states, including two CD8+ T cell-related signatures that potentially reflect pre-exhaustion and exhaustion states, cellular ecosystem dynamics, and the functional characteristics that accompany or drive PC progression.

### Myeloid cell subtypes exhibit characteristics supportive of PC progression

Recent pan-cancer studies have delineated the heterogeneity and functional properties of myeloid cells in the TME of diverse tumor types^75, 76, 77, 78^. However, these studies did not include PC samples, leaving an incomplete understanding of the myeloid composition in PC. Overall, 77,685 cells were classified as myeloid with subclassification into six major cell types and 13 cell states^72, 75, 76^ (Figures S19A, S19B, 6A, and 6B). These included a mast cell cluster, two monocyte clusters, four macrophage cluster, three dendritic cell (DC) clusters, a neutrophil cluster, and a cycling myeloid cluster. Notably, macrophages were the predominant population among myeloid cells and were associated with an unfavorable prognosis (Figures S19C and S19D). Moreover, using curated myeloid cell associated pathway signatures from recent publications^72, 75^ (Table S3), we identified two pathway modules that were enriched in advanced PC, including macrophage functions (e.g., migration, M2 polarization, phagocytosis, and tissue resident) and metabolism-related pathways (e.g., glycolysis, fatty acid metabolism, and oxidative phosphorylation) (Figure S19E), suggesting that cancer cells may modify macrophage phenotypes during PC progression.

We further examined the tissue preference and distribution of all myeloid cell subtypes across a trajectory of PC progression. We quantitated higher proportions of Mph_SPP1 cells in tumor tissues relative to benign prostate tissues and a significant enrichment of these cells in advanced PC (Figures S19F, 6C, and 6D), indicating that this subcluster accumulates in tumors and may contribute to tumorigenesis and tumor progression^72, 75^. Furthermore, the abundance of Mph_SPP1 cells increased in lymphatic micro-metastases^18^, and further overt PC lymphatic metastases (Figure 6E). This suggests that alterations in immune cell gene expression within lymph nodes (LNs) might precede the onset of actual metastasis and progression, potentially contributing to the establishment of a pre-metastatic niche. Using independent scRNA-seq data and bulk RNA-seq data, we confirmed the accumulation of Mph_SPP1 cells during PC pathological progression (Figures 6F, 6G, and S19G).

**Figure 6.**
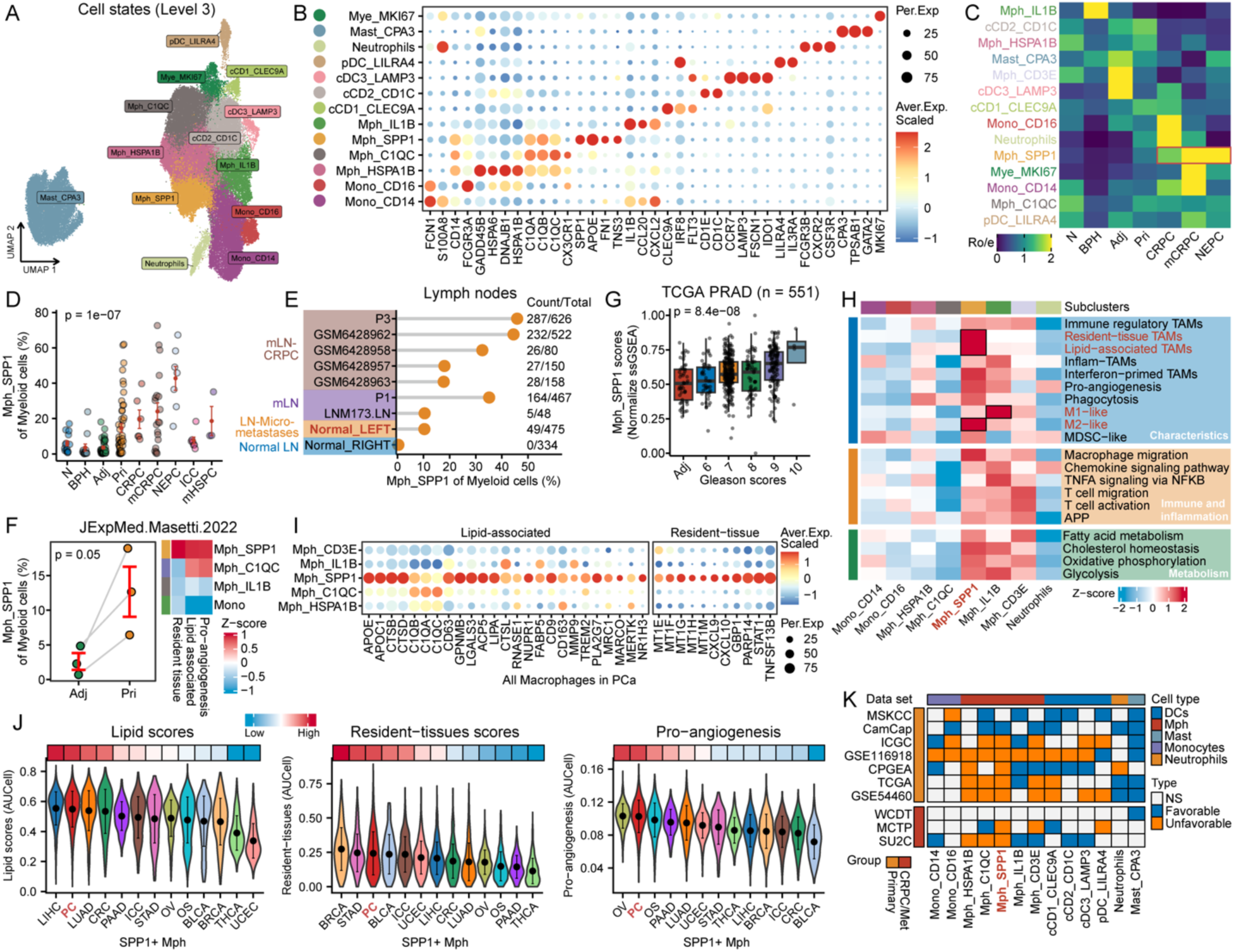
Characteristics of myeloid cells during PC progression. A. UMAP plot of all myeloid cells, colored by cell states. Mono, monocytes; Mph, macrophages; DC, dendritic cells. B. Dot plot showing representative marker genes for each cell cluster. C. Tissue preference of each myeloid cluster estimated by Ro/e score. D. Scatter diagram showing the frequency change of Mph_SPP1 with PC progression. E. Lollipop chart showing the distribution of Mph_SPP1 in lymphatic tissues across various progression stages of PC. F. The composite plots showing the validation results of frequency (left) and pathway signaling activities (right) for Mph_SPP1 in an independent PC scRNA-seq dataset. G. Boxplot showing the distribution of Mph_SPP1 gene signature scores among patients with different Gleason scores in the TCGA PRAD cohort. H. Heatmap illustrating expression of 20 curated gene signatures across cell clusters of monocytes, macrophages and neutrophils. I. The expression of representative marker genes associated with lipid metabolism and resident tissues for each macrophage cluster. J. Violin plots showing the comparison of Mph_SPP1 in terms of lipid activity, tissue residency, and pro-angiogenesis between PC and other cancer types. Cancer types are arranged based on their respective pathway activities. K. Univariate Cox regression analysis for each myeloid cluster. Survival index using BCR for primary PC and OS for CRPC/Met PC.

We next delineated the inferred functions and characteristics for each monocyte, macrophage, or DC cluster based on their gene expression programs (Figures 6H, 6I, S20A, and S20B). Notably, Mph_SPP1 cells exhibited heightened pro-angiogenesis, M2 polarization, and diminished M1 polarization related-signature activity (Figure 6H). Interestingly, we noted a significant enrichment of tissue resident and lipid characteristics in Mph_SPP1 of PC (Figures 6H and 6I). Further pan-cancer analysis revealed that Mph_SPP1 cells from PC exhibited the highest levels of lipid, tissue resident, and pro-angiogenesis features, compared to Mph_SPP1 cells found in other cancer types (Figures 6J and S19H). Notably, we also determined that the signature genes of Mph_SPP1 cells were correlated with unfavorable prognosis in multiple PC cohorts (Figure 6K). Collectively, these data support the potential for myeloid cell subtypes to influence PC behavior, unveiling distinct characteristics of Mph_SPP1 cells and emphasizing their potential role in tumor progression and metastasis.

### Co-localization of mCAFs and Mph_SPP1 in PC is associated with worse patient survival

To evaluate the potential contribution of cellular spatial relationships toward PC pathobiology, we assessed how the presence and proximity of one cell type associated with specific cell phenotypes of other lineages. Correlation analysis of subpopulation proportions identified potential interaction networks of major cell types and cell states (Figures S21A and S21B). Notably, a significant correlation was observed between the subpopulation proportions of mCAFs and Mph_SPP1 (Figures S21A and S21B). We also ranked the subpopulation proportion correlation between Mph_SPP1 and each of the other cell states (Figure 7A), which revealed that mCAFs were the one of the cell types most correlated with Mph_SPP1 cells. The subpopulation proportions of Treg_FOXP3 and CD8Tex_terminal T cells also exhibited a strong correlation with Mph_SPP1. Mph_SPP1 also occupied the top position in the correlation ranking of mCAFs (Figure S22A). The associations between the Mph_SPP1 signature and other gene signatures, including mCAFs, Tregs, and T cell exhaustion, were validated across 12 independent PC cohorts (Figure S22B).

**Figure 7.**
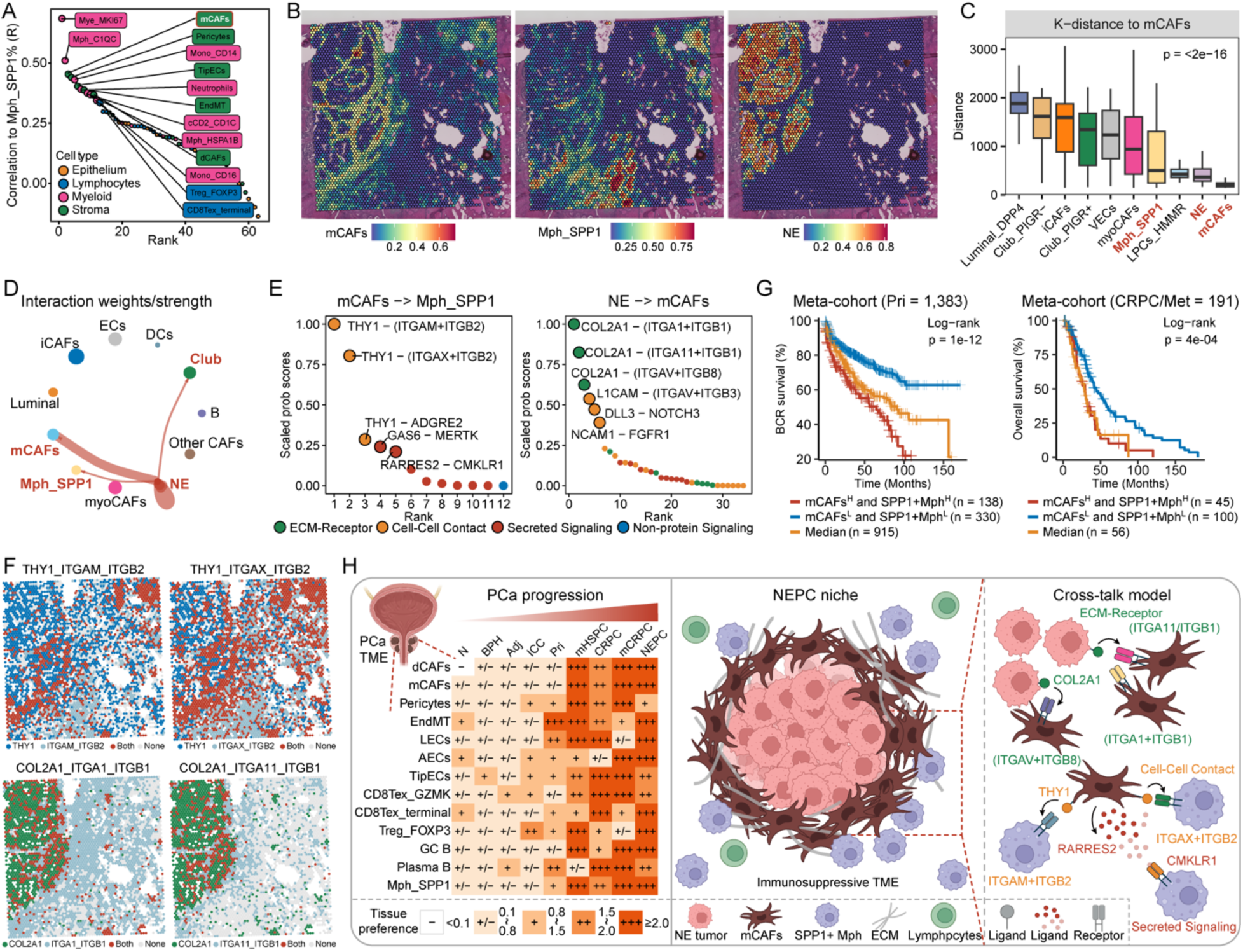
Co-localization of mCAFs and Mph_SPP1 localized as a tumor immune barrier that is associated with worse patient survival. A. Spearman correlation analyses between the cell proportion of Mph_SPP1 and other cell subpopulations. Each data point on the scatter plot represents a cell subpopulation, ordered by its correlation to Mph_SPP1. B. The spatial distribution of mCAFs (left), Mph_SPP1 (middle) and NE tumor cells (right), as inferred by the RCTD deconvolution algorithm. C. Box plot showing RCTD-based spatial K-distance of mCAFs to other cell subpopulations. D. Circle plot of the cellular crosstalk of NE cells toward the other cell subclusters. Dot size and edge width indicated the communication probability. E. Scatter plots showing potential ligand-receptor gene pairs involved the interaction between mCAFs and Mph_SPP1 (left), or well as NE and mCAFs (right). Each scatter point represents a ligand-receptor pair, ordered by its normalized communication probability. F. The feature plots showing the expression distribution of each ligand-receptor gene pair by binarizing their expression in each spot. Each spot is colored based on whether it only expresses the ligand, the receptor or both signaling molecules. G. Kaplan-Meier survival curves for BCR (for primary PC, left) and OS (for CRPC/Met PC, right) based on both mCAFs and Mph_SPP1 signature scores in the two PC meta-cohorts. H. Graphical summary of advanced PC TME niche and the cell-cell interactions among mCAFs, Mph_SPP1 and NE tumor cells. +++, Ro/e >=2; ++, 1.5 < Ro/e < 2; +, 1 < Ro/e < 1.5; +/−, 0.1 < Ro/e < 1; −, 0 < Ro/e < 0.1. N, normal; BPH, benign prostate hyperplasia; Adj, normal tissues adjacent to the tumors. ECM, extracellular matrix.

To further assess the spatial organization of Mph_SPP1 and mCAFs, we evaluated a stRNA-seq dataset derived from localized PC using the robust cell type decomposition (RCTD) method^79^. We deconvoluted each spot in the stRNA-seq tissue section using our PCCAT data as a reference (Figures S22C and S22D). The results validated the inferred spatial proximity between Mph_SPP1 and mCAFs identified through scRNA-seq data and further demonstrated that mCAFs were distinctly co-located adjacent to NE tumor cells (Figures 7B, 7C, and S22E). Moreover, the signature score of Mph_SPP1 exhibited strong positive correlation with the signature scores of mCAFs, T cells exhibiting a Treg, and T cell exhaustion state in multiple spatial slices (Figure S23A), indicating the presence of an immunosuppressive niche that may provide a favorable environment for tumor growth.

To further explore the potential crosstalk among NE cells, mCAFs, and Mph_SPP1, we identified the top ligand/receptor genes for each cell cluster and the enriched GO terms that may contribute to their intercellular communication (Figure S23B, S23C, and S24A). We also used the CellChat2 method^80^ to elucidate spatial cell-to-cell communication patterns, revealing a notable interaction between NE tumor cells and mCAFs, Mph_SPP1, as well as Club cells (Figures 7D and S23D). Moreover, we identified a subset of ligand/receptor gene pairs indicating potential communication mechanisms between Mph_SPP1 and mCAFs by overlapping the top signaling pairs at both the single-cell and spatial levels (Figures 7E, 7F, S23E, and S23F). The potential interactions derived from these analyses include THY1_ITGAM_ITGB2, THY1_ITGAX_ITGB2, and RARRES2_CMKLR1. In addition, we found that patients with both high Mph_SPP1 and mCAFs exhibited the shortest survival compared to other groups (Figures 7G), supporting the hypothesis that these two cell types can synergistically promote tumor progression. Lastly, we explored the potential interactions between NE tumor cells and mCAFs, identifying the top ligand-receptor gene pairs (Figures 7E, S24B, and S24C), including COL2A1-ITGA1_ITGB1, L1CAM-ITGAV_ITGB3, and DDL3-NOTCH3. Interestingly, these findings were not limited to NEPC, but were also observed in a tumor histologically classified as ‘invasive’ in a section of PC (Figures S24D-S24F). High signature scores of NE cells and mCAFs were also both associated with enzalutamide resistance (Figure S24G). Taken together, these data present a novel concept suggesting that the co-localization and consequent interactions between mCAFs and Mph_SPP1 may constitute an immune barrier in advanced PC linked to poor patient survival (Figure 7H).

## Discussion

Localized and metastatic PC comprise complex and dynamic ecosystems that influence key aspects of pathology spanning cancer initiation through responses to therapeutics. Understanding the composition and cellular relationships existing in these TMEs is now enabled through detailed assessments of cell phenotypes using single cell transcriptome analyses. Although several single-cell analyses have been conducted on PC, most have involved small sample sizes and tended to concentrate on a specific stage of PC. Consequently, an integrated view of disease progression and responses to treatments is lacking^81^. To fill in this gap, we produced a large-scale atlas of single-cell transcriptomes of PC through the integration of 19 datasets comprising over 710,000 cells from 197 samples spanning various disease stages. Based on this large-scale atlas, we characterized the dynamic cellular ecosystem, determined functional characteristics, and evaluated the clinical relevance of diverse cell states in the TME during tumorigenesis and PC progression to metastasis.

To address potential dataset-specific batch effects while retaining biological information, we combined pre-annotation information and data integration algorithms to construct the PC reference atlas named PCCAT. Interrogation of this atlas provides a high-resolution view of the PC TME, delineating 7 major cell types (Level 1), 23 refining cell subtypes (Level 2) and 70 transitional cell states (Level 3). Notably, a significant challenge in compiling distinct scRNA-seq datasets arises from varied definitions of cell types used by different research groups, leading to inconsistencies in naming schemas^38, 82^. To address this, we developed an interactive web portal and an automated alignment tool (https://pccat.net) to achieve a consensus cell type re-annotation classifier along with corresponding marker genes.

As predominant nonmalignant epithelial cells, the characteristics of the basal, hillock and club cell types have been defined previously in the normal prostate^36^. Analyses of these cell types suggest that these nonmalignant cells may undergo a loss of their original cell identity and acquire luminal-like states during PC progression. More importantly, we determined that club cells may contribute to tumor progression beyond previously reported roles in carcinogenesis^30, 44^. With respect to malignant epithelial cell types, tumor cells classified as Lum_DPP4 and Lum_KLK4 exhibited features associated with low malignancy rate, while Lum_PCA3 and Lum_ERG cells exhibit features predominantly associated with malignant behavior. Additionally, we identified NE-like tumor cell states associated with poor prognosis, including LPCs_HMMR and three classic NE subpopulations enriched in advanced PC. Furthermore, the integration of bulk RNA-seq data provided valuable insights into the correlation between epithelial subpopulations, genotypic features, and clinical outcomes. These findings could be exploited to derive rationale for potential prognostic biomarkers and personalized therapeutic combination strategies. For example, our study unveiled insights into treatment outcomes, highlighting associations between specific subpopulations (e.g., NE_FOXA2) and distinctive features (such as high MHC expression, low AR activity, and stemness epithelial program) linked to enzalutamide resistance. Enzalutamide resistance gene signatures were further developed, demonstrating robust predictive capability for therapeutic outcomes. Taken together, these results highlight the diversity of neoplastic epithelial cells in the prostate and delineate characteristics that are associated with PC progression and therapeutic resistance.

Stromal cells, including endothelial cells and fibroblasts, are recognized as vital components of the TME that contribute to tumor pathophysiology and importantly to prognosis and response to therapy. However, our understanding of endothelial cells at the single-cell level is currently limited^45, 61, 81^. The in-depth analysis of endothelial cells in various stages of PC identified alterations in immune-related pathways in localized PC and key signaling pathways in advanced PC (e.g., Myc targets, Notch signaling, and angiogenesis), suggesting dynamic changes in endothelial cell function during cancer progression. Additionally, time-series analysis identified gene clusters with distinct expression patterns and functions in endothelial cells throughout PC progression, emphasizing the complexity of their role in the disease. Recent studies suggested that the microenvironment surrounding the tumor, in addition to tumor cell intrinsic features, is essential for understanding recurrence and progression^66^. Hence, we delved into the enrichment patterns of several stromal cell clusters (e.g., TipECs, mCAFs, and pericytes) in advanced PC, revealing their association with unfavorable outcomes. Notably, we observed that a small subset of phenotypically distinct fibroblasts, hypothesized to originate from adjacent normal fibroblasts, were associated with tumor recurrence.

Previous studies have been constrained by a limited number of TME cells, attributed to the small size of PC samples and immunologically ’cold’ characteristics^81^. Here, leveraging the largest scRNA-seq atlas of PC, we identified and tracked diverse lymphocyte and myeloid cell states throughout PC progression. Notably, we detected the co-enrichment of several cell states within the immunosuppressive TME of advanced PC, featuring activated Treg cells, exhausted CD8+ T cells, and SPP1-expressing macrophages. Additionally, we mapped the evolutionary trajectory from CD8Tn_CCR7 to effector memory CD8+ T cells or exhausted CD8+ T cells, identifying dynamic molecular changes during this transition. These changes include aberrant interferon signaling, TCR pathways, and PD-1 pathways, consistent with recent reports^26^. Finally, we developed and validated an exhausted T cell-related immunotherapy response signature across multiple published datasets. These findings provide valuable insights that may enhance cancer immunotherapy strategies for PC and potentially other cancers.

The interactions between CAFs and tumor cells, as well as tumor-infiltrating immune cells, have been identified as key factors in promoting tumor progression^45^. Here, we observed a significant correlation between the subpopulation proportions of mCAFs and Mph_SPP1. Spatial analysis further confirmed the co-localization and provided evidence for active communication between these two cell types. This observation aligns with previous study in colorectal cancer, which demonstrated a strong correlation and spatial co-localization of FAP+ CAFs and SPP1+ macrophages within tumor tissues, which can facilitate the formation of desmoplastic structures and synergistically promote tumor progression^83^. Moreover, we determined that mCAFs exist in close proximity to, and envelop NE tumor cells, with evidence for inter-cellular communication. The interaction patterns among tumor cells, mCAFs, and Mph_SPP1 imply a complex interplay, forming a tumor immune barrier associated with poor patient survival in advanced PC. Overall, this study provides a deep view into the cellular and molecular characteristics of PC, from early indolent disease through tumorigenesis and metastasis. These findings held the potential to advance PC diagnosis, refine patient stratification, and inform future research aimed at developing therapeutic strategies that address the complexity of these tumors and their microenvironments.

## Methods

### Single-cell RNA-seq data collection and processing

In this study, we collected a total of 19 previously published human PC scRNA-seq datasets as reference core atlas (PCCAT) from 197 samples covering various tumor stages (Table S1). In addition, we incorporated two published based on fluorescence-activated cell sorting (FACS) for validation purposes. As previously described^39, 41^, we first performed quality control, pre-annotation and doublets detection for each dataset. Low quality cells (defined as cells with fewer than 250 detected genes, fewer than 500 transcripts, or more than 20% mitochondrial content), erythroid cells, and hematopoietic stem and progenitor cells were excluded to mitigate potential interference with the further analyses. High dropout genes (defined as those expressed in fewer than 500 cells) were removed to avoid interference with subsequent analyses. Besides, to avoid unexpected noise and expression artefacts by dissociation, genes associated with mitochondria and ribosome were excluded. After pre-annotation and merging, 716,763 high-quality cells with 20,560 genes from 197 samples were preserved for subsequent analyses. To address potential batch effects stemming from diverse platforms and protocols, we employed 11 classical integration algorithms (including BBKNN, CCA, RPCA, LIGER, Harmony, fastMNN, scVI, scANVI, Scanorama, Connos, and Combat) alongside an unintegrated method as a reference. Clustering and single-cell distribution were visualized using Uniform Manifold Approximation and Projection (UMAP) with the Leiden algorithm. Based on pre-annotation labels and scib evaluation^84^, we determined that the BBKNN method was more suitable for our PCCAT and was thus chosen for further analyses. Subsequently, cell clusters were annotated based on previously reported marker genes^39, 41^ and the combined automatic annotation method Celltypist^82^. During cell subclustering analyses, a similar procedure was applied, and a second round of quality control was performed to exclude clusters with low quality and high doublet scores.

### Spatial transcriptomic data collection and processing

stRNA-seq datasets were downloaded from the Gene Expression Omnibus (GEO) or 10X Genomics database (see Table S1). For GSE230282, pathologists have annotated this tissue as NEPC coexisting with HSPC^27^. Each stRNA-seq data was standardized and corrected using the Sctransform method, following the corresponding Seurat tutorial. The RCTD method^79^ was employed to deconvolute each spot in the 10X Visium or Slide-seqV2 slices using scRNA-seq data as a reference (see Supplementary Methods). For the rough annotation, we assigned each spot to a specific cell type as the first class based on the highest probabilistic proportion.

### Bulk dataset selection and preparation

In this study, we assembled over 3000 samples from publicly available and in-house human bulk PC datasets^3^ (Table S1). As previously described^39, 41^, the RNA-seq datasets including TCGA PRAD^7^, ICGC PRAD, SU2C^5^, MCTP^8^, WCDT^9^, CPGEA^10^, PCaProfiler^11^, and SciTranslMed.Subudhi.2020^85^ were download according to corresponding publication. The microarray data including MSKCC^13^, CamCap^14^, GSE54460^15^, and GSE116918^16^ were acquired from GEO. Besides, we gathered transcriptomic data of PC cell lines from the GEO database (Table S1). For all RNA-seq data, Transcript Per Million (TPM) value was calculated.

### Identification of malignant cells

To distinguish the malignant cells from non-malignant cells, interCNVpy (v0.4.3) was employed to infer large-scale chromosomal copy number variations, following the corresponding tutorial. We used cnv.tl.infercnv function to compute rolling average gene expression changes with parameter ‘window_size=100’ for each scRNA-seq dataset. Lastly, classical tumor markers and top 50 up-regulated DEGs between tumor and paired non-tumor samples from TCGA-PRAD were applied to validate the identified malignant and non-malignant cells (Figures S5E and S5F).

### Gene regulatory network analysis

For the gene regulatory network analysis, we used SCENIC algorithm^62^ in the Python (pySCENIC) to inference the regulatory networks from the metacells following the instructions available online. TFs were ranked according to the specificity scores of the regulons by combining the activity score of the regulator and meta-clusters.

### Gene set enrichment analyses

As previously described^39, 41^, we conducted GO, Reactome, and rank-based gene set enrichment analysis (GSEA) using the R package clusterProfiler (v4.7.1). For bulk transcriptomic data, we employed the single sample GSEA (ssGSEA) method with the R package GSVA. For scRNA-seq and stRNA-seq data, AUCell algorithm^62^ was employed to mitigate the impact of dropouts and inherent technical variability.

### Statistical analysis

Statistical analyses were performed using the R (v.4.1.0) platform, including Student’s t-test, Wilcoxon rank-sum test, one-way ANOVA, Kruskal-Wallis tests, Pearson’s correlation, and Spearman’s correlation. The Benjamini-Hochberg (BH) method was applied to estimate the false discovery rate for multiple testing. The tissue samples from patients used in the survival analyses were divided into distinct groups based on the quantile expression of the proposed gene signature using the surv_cutpoint function of the survminer R package. Subsequently, Kaplan-Meier analysis with log-rank tests was conducted to assess survival differences between the groups. The proportional hazards assumption for the COX model was validated for all variables using the Schoenfeld residuals test. A p-value < 0.05 was considered statistically significant with two-sided testing.

## Acknowledgments

This work was supported by the following funding: Department of Defense Prostate Cancer Idea Development Award W81XWH2110539, Prostate Cancer Data Science Award HT94252410551, NIH R01GM147365 and the Silver Family Innovation Foundation Award (to Z.X.). NIH P50CA097186, NIH R01CA234715, NIH R01CA266452 and the Prostate Cancer Foundation (to P.S.N.). Breast Cancer Research Foundation and NIH U01CA253472 and U01CA217842 (to G.B.M.). NIH R37CA263592 (to A.E.M.). NIH R01CA251245, NIH R01CA282005, NCCN/Pfizer/Astellas Award, Joint Institute for Cancer Research Award, and Prostate Cancer Foundation (to J.J.A.). The content is solely the responsibility of the authors and does not necessarily represent the official views of the funders.

## Author contributions

F.Z. and Z.X. conceived the idea. F.Z., J.Z., C.C. and T.Z. performed computational analyses. X.Z. developed the website. G.V.T., R.C.S., J.J.A., A.E.M., G.B.M. and P.S.N. interpreted the data and provided clinical insights. Z.X. and P.S.N. jointly supervised the study. F.Z., P.S.N. and Z.X. wrote the manuscript. All other authors provided critical feedback and approved the final manuscript.

## Data and code availability

This paper analyzes existing, publicly available data. These accession numbers for the datasets are listed in Table S1. Processed scRNA-seq data has been deposited on PCCAT (https://pccat.net). All original code has been deposited on GitHub (https://github.com/Famingzhao/PCCAT).

## Supplemental material

**Figure S1-S24.**

**Supplemental table 1.** Metadata of scRNA-seq and bulk RNA-seq cohorts used in this study.

**Supplemental table 2.** Gene signatures generated by this study.

**Supplemental table 3.** Gene sets curated in this study.

**Supplemental table 4.** Gene lists for four clusters of CD8+ T cells identified through pseudotime analysis.

