## Supplementary for "Deciphering single-cell heterogeneity and cellular ecosystem dynamics during prostate cancer progression"

### Supplementary Figures

#### Figure S1

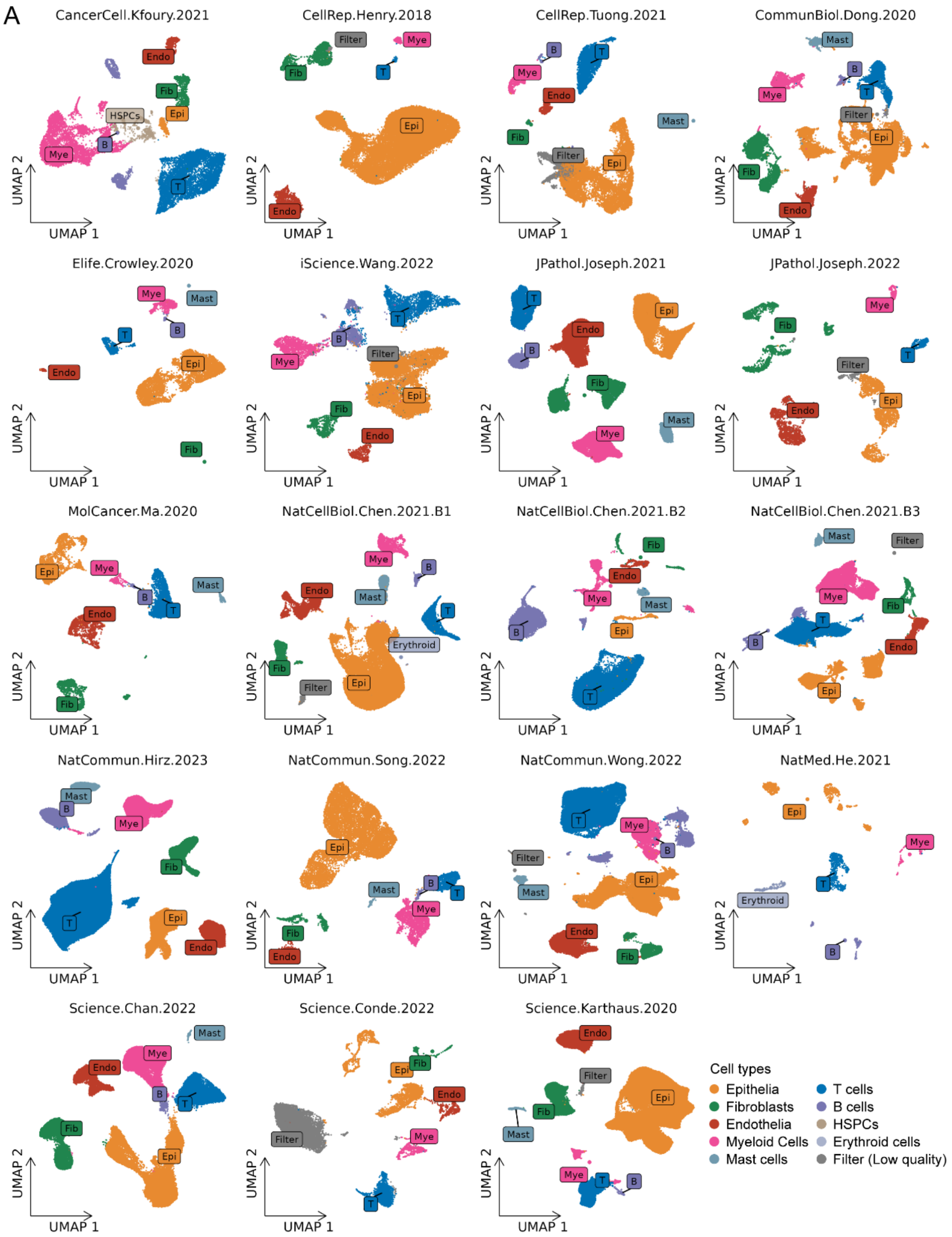


**Figure S1. UMAP plots depicting pre-annotated cell cluster across the 19 independent datasets, related to Figure 1.**

**Figure S2**

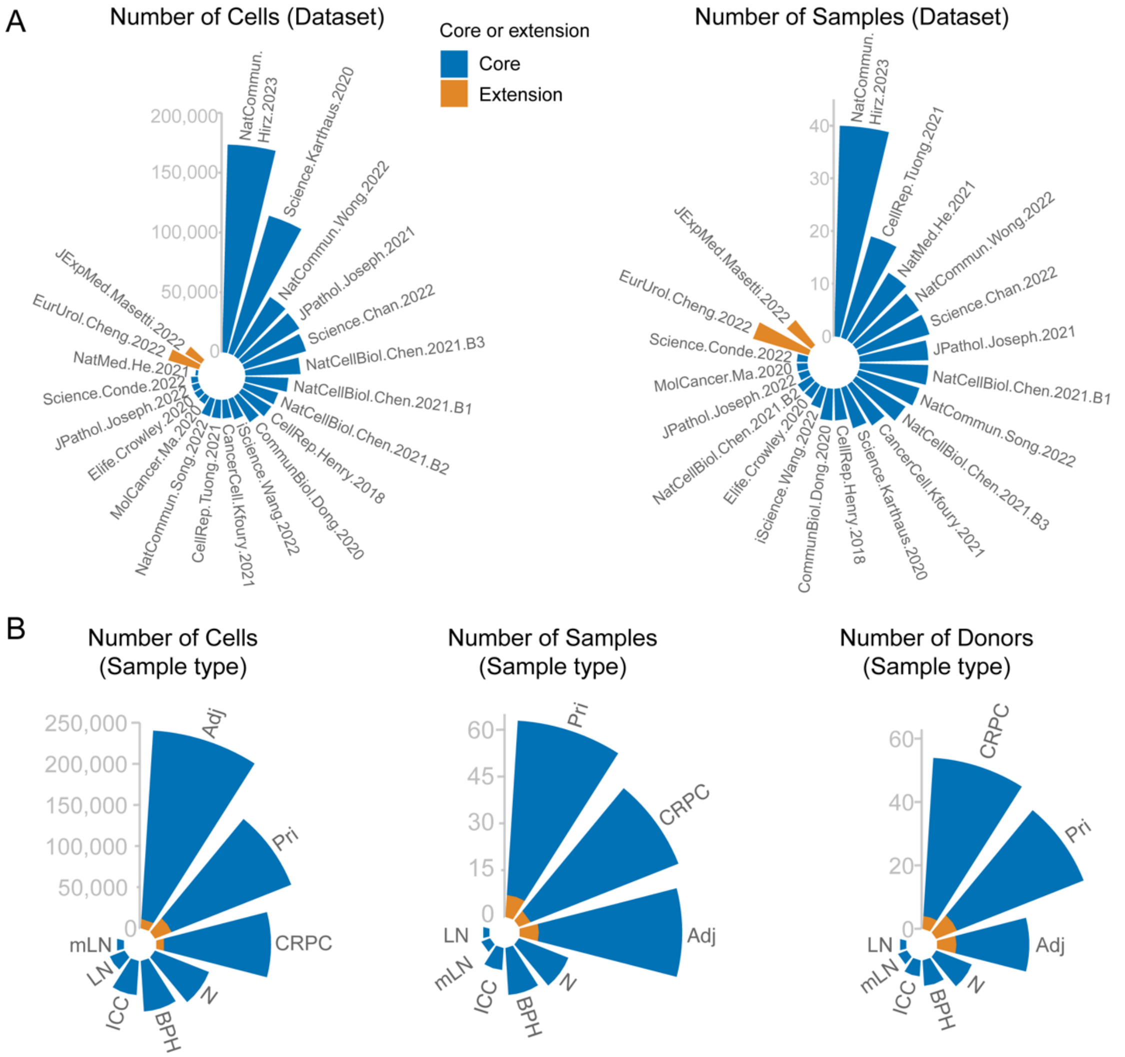


**Figure S2. Cell and sample composition of the PCCAT, related to Figure 1.**

A and B. Nightingale Rose plots showing the number of cells, samples, or donors of data from different datasets (A), and disease groups (B) of PC.

**Figure S3**

| Method | Bio conservation | | | | | Batch correction | | | | Aggregate score | | |
|---|---|---|---|---|---|---|---|---|---|---|---|---|
| | Isolated labels | KMeans NMI | KMeans ARI | Silhouette label | cLISI | Silhouette batch | iLISI | KBET | Graph connectivity | Batch correction | Bio conservation | Total |
| **BBKNN** | 0.44 | 0.85 | 0.87 | 0.71 | 1.00 | 0.74 | 0.06 | 0.02 | 0.99 | 0.45 | 0.77 | 0.65 |
| **CCA** | 0.39 | 0.82 | 0.71 | 0.72 | 1.00 | 0.72 | 0.16 | 0.28 | 0.98 | 0.54 | 0.73 | 0.65 |
| **RPCA** | 0.64 | 0.72 | 0.53 | 0.74 | 1.00 | 0.73 | 0.15 | 0.24 | 0.98 | 0.53 | 0.73 | 0.65 |
| **LIGER** | 0.59 | 0.57 | 0.41 | 0.64 | 1.00 | 0.84 | 0.19 | 0.52 | 0.96 | 0.63 | 0.64 | 0.64 |
| **Harmony** | 0.40 | 0.72 | 0.60 | 0.66 | 1.00 | 0.78 | 0.15 | 0.37 | 0.99 | 0.58 | 0.67 | 0.63 |
| **fastMNN** | 0.58 | 0.72 | 0.56 | 0.74 | 1.00 | 0.69 | 0.10 | 0.07 | 0.96 | 0.46 | 0.72 | 0.61 |
| **scVI** | 0.59 | 0.69 | 0.56 | 0.67 | 1.00 | 0.67 | 0.06 | 0.03 | 0.96 | 0.43 | 0.70 | 0.59 |
| **scANVI** | 0.57 | 0.63 | 0.50 | 0.65 | 1.00 | 0.70 | 0.08 | 0.05 | 0.96 | 0.45 | 0.67 | 0.58 |
| **Scanorama** | 0.57 | 0.61 | 0.46 | 0.63 | 1.00 | 0.73 | 0.02 | 0.00 | 0.98 | 0.43 | 0.65 | 0.57 |
| **Unintegrated** | 0.55 | 0.54 | 0.40 | 0.56 | 1.00 | 0.65 | 0.00 | 0.00 | 0.82 | 0.37 | 0.61 | 0.51 |
| **Conos** | 0.50 | 0.31 | 0.15 | 0.50 | 0.95 | 0.66 | 0.22 | 0.30 | 0.76 | 0.48 | 0.48 | 0.48 |
| **Combat** | 0.55 | 0.13 | 0.07 | 0.43 | 0.97 | 0.64 | 0.00 | 0.00 | 0.51 | 0.29 | 0.43 | 0.37 |

**Figure S3. Summary of the dataset integration benchmarking results for PC scRNA-seq datasets, related to Figure 1.**

#### Figure S4

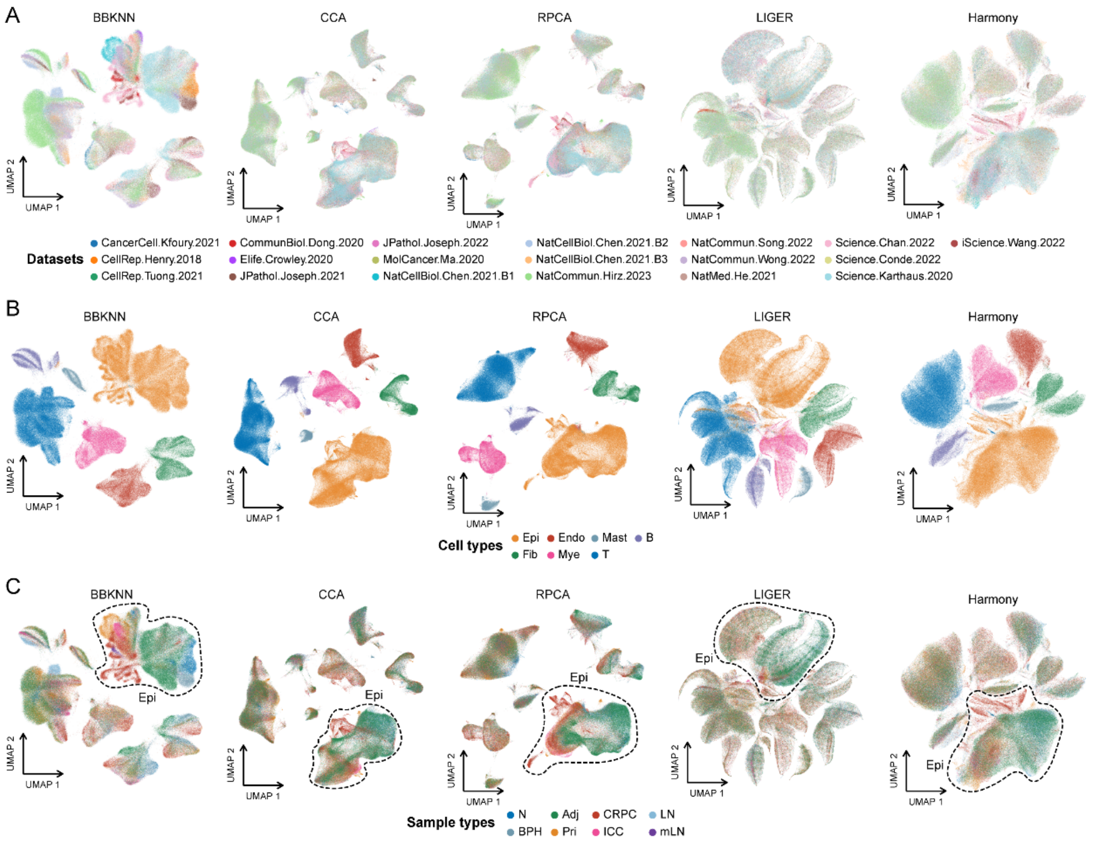


**Figure S4. UMAP plots of the dataset integration benchmarking results for PC scRNA-seq datasets, related to Figure 1.**

A-C. UMAP plots depicting 716,763 cells from 241 human samples in core PC scRNA-seq altlas, colored by different datasets (A), pre-annotated major cell types (B), or disease groups (C).

**Figure S5**

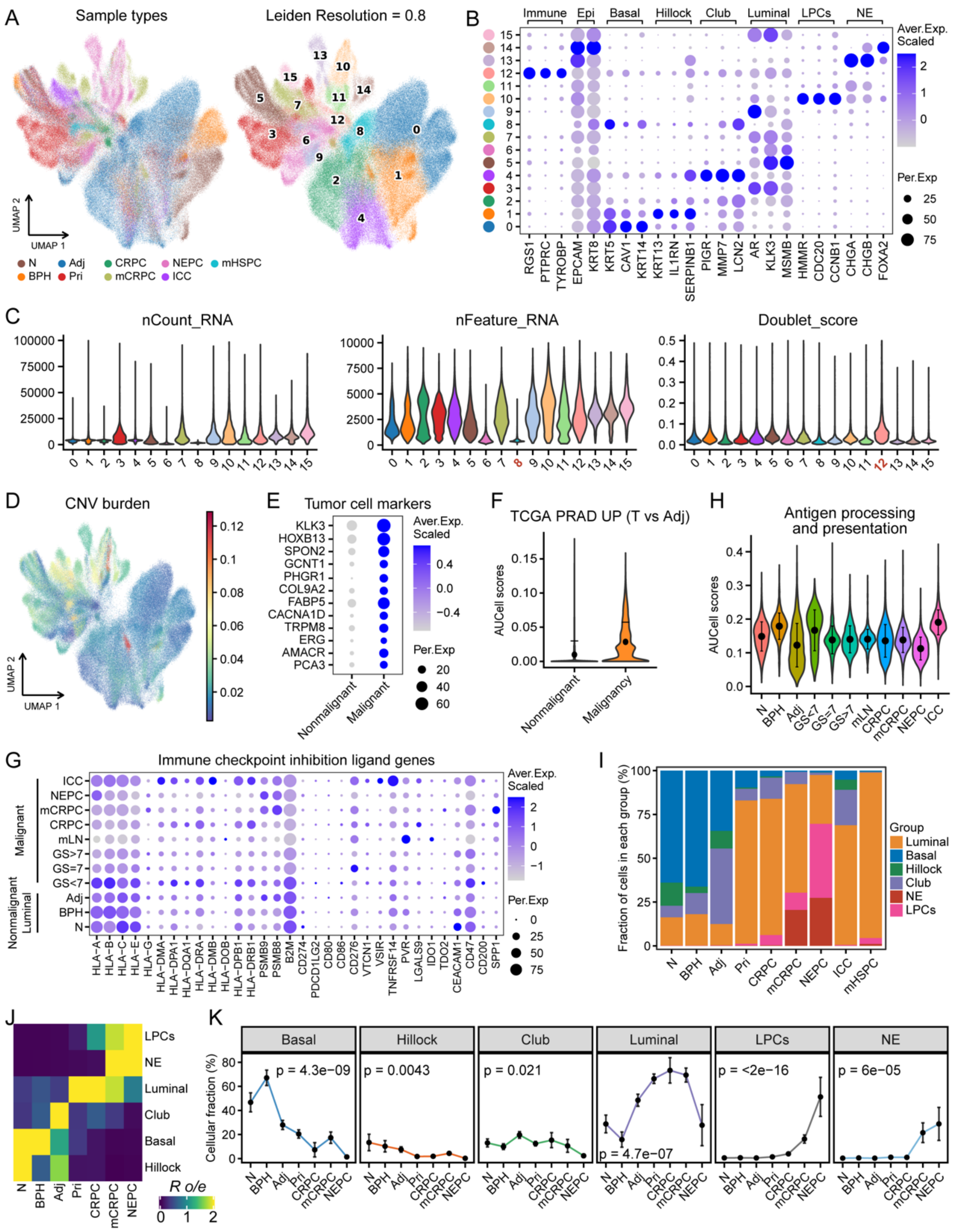


**Figure S5. Re-cluster analysis of epithelial cell subsets, related to Figure 2.**

A. UMAP plots showing 285,850 individual epithelial cells that colored by different disease groups (left) and cell clusters (right).

B. Dot plot of representative marker genes for each cluster.

C. Violin plots showing the quality control (QC) index for each cell cluster, including nCount_RNA (left), nFeature_RNA (middle), and doublet scores (right) inferred by Scrublet.

D. UMAP plots showing the distribution of CNV scores inferred by using inferCNVpy.

E. Dot plot showing the expression of classical marker genes associated with malignant cells in PC.

F. AUCell enrichment analysis comparing the activity of tumor versus paired adjacent normal signature (top 50 up-regulated DEGs) between inferred nonmalignant and malignant cells.

G. The features of immune checkpoint inhibition ligand genes expressed in different stages of PC nonmalignant and malignant cells.

H. Violin plots showing the distribution of antigen processing and presentation (APP) pathway scores in nonmalignant and malignant cells across various disease groups.

I. Bar plots showing the proportion of different epithelial cell types in each disease group.

J. Tissue preference of each epithelial type estimated by Ro/e score.

K. Dynamic changes in cellular fraction for each major epithelial cell type during PC progression.

**Figure S6**

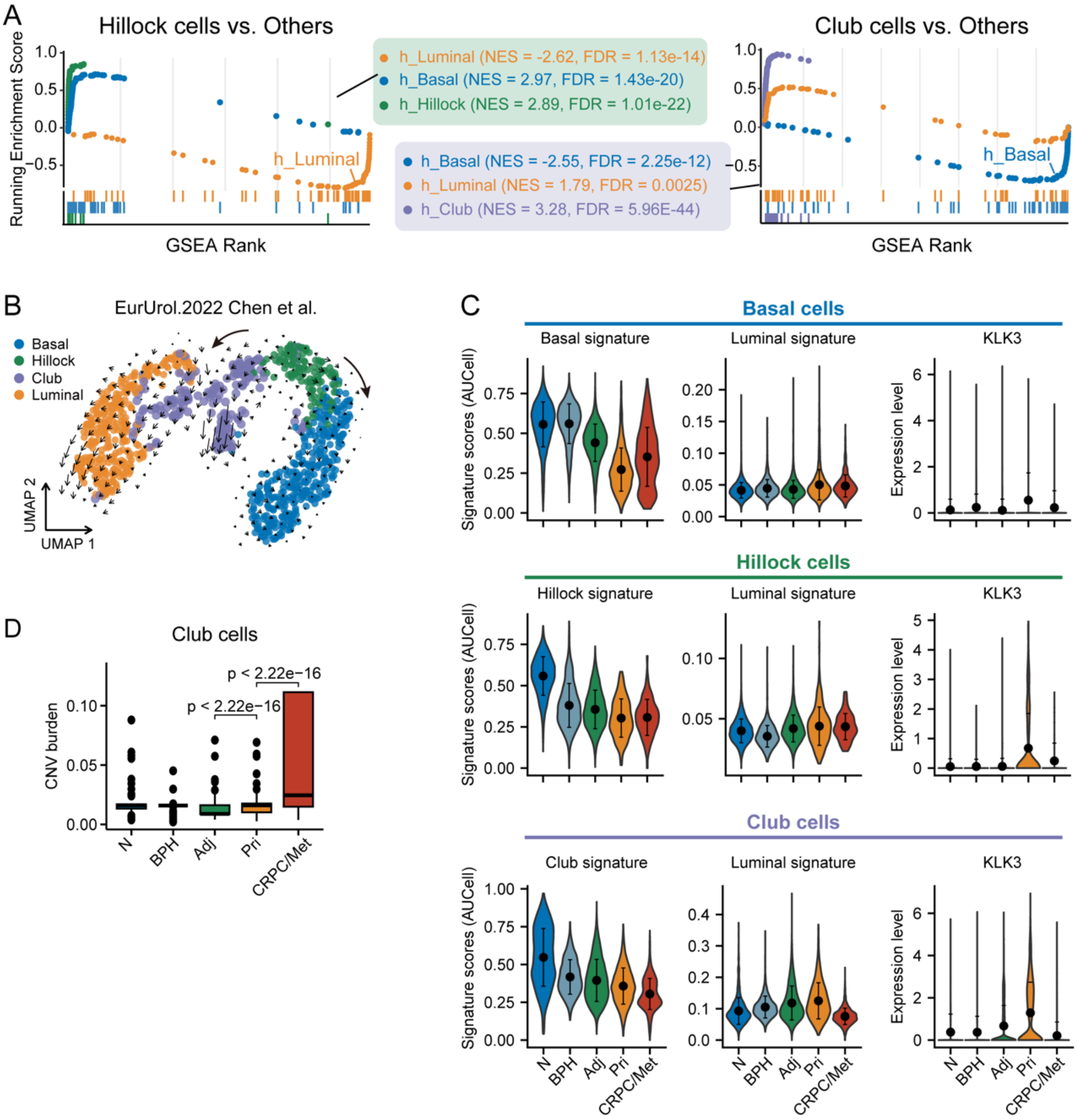


**Figure S6. Basic characteristics of epithelial cell subsets during PC progression, related to Figure 2.**

A. GSEA plots showing the dynamic changes of epithelial features in hillock cells versus other epithelial cells (left), or club cells versus other epithelial cells (right).

B. RNA velocities overlaid on UMAP showing major state transition paths from club cells to Luminal cells, as well as hillock cells to basal cells. Arrows on a grid show the RNA velocity field, and dots are colored by cell types.

C. AUCell enrichment analysis showing the dynamic changes of epithelial feature signature scores or *KLK3* expression during PC progression.

D. Box plot showing the distribution of the CNV scores among different groups within Club cells.

**Figure S7**

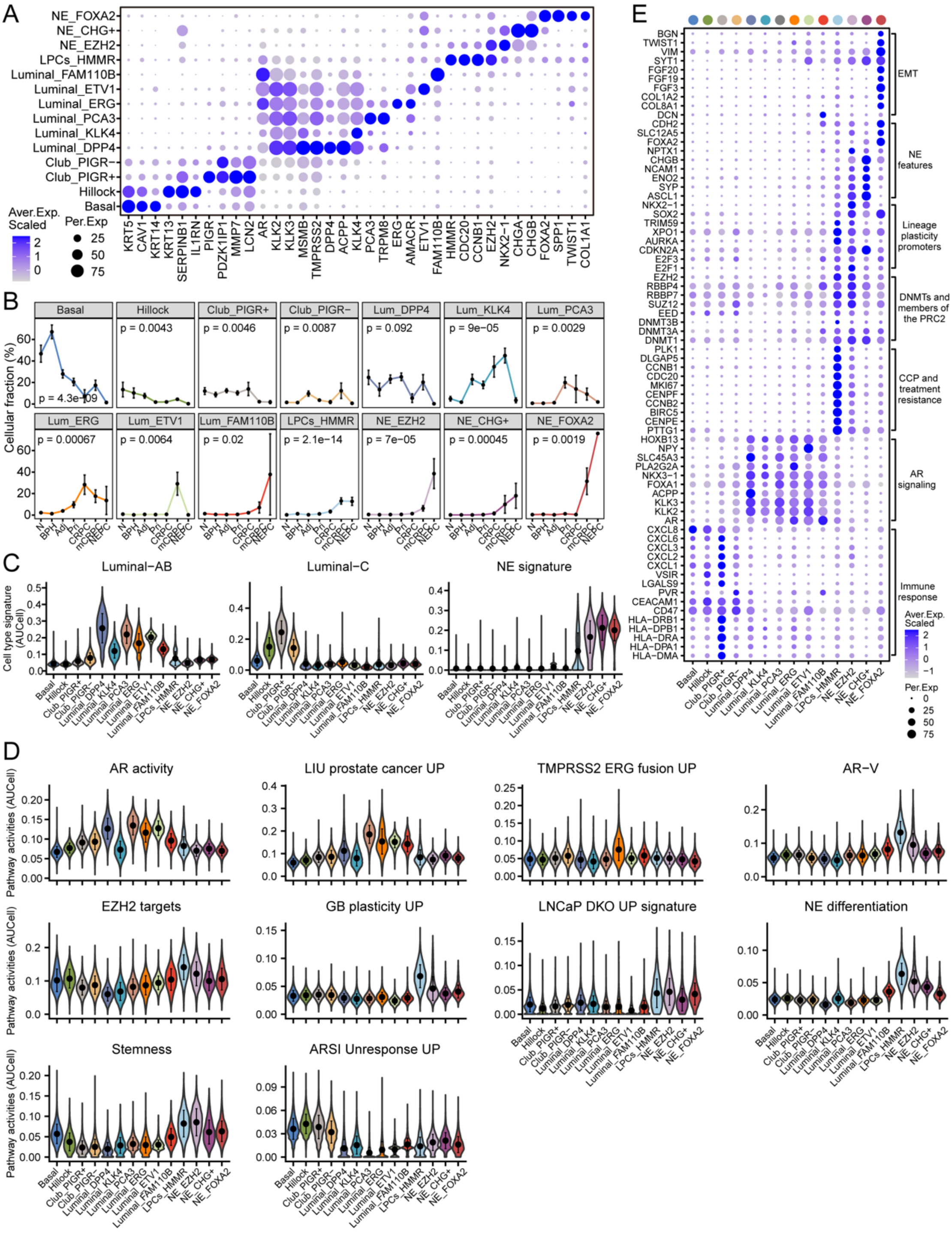


**Figure S7. Molecular characteristics of different epithelial cell subpopulations during PC progression, related to Figure 2.**

A. Dot plot of representative marker genes for each epithelial cell state.

B. Dynamic changes in cellular fraction for each epithelial cell cluster during PC progression.

C and D. AUCell enrichment analysis showing distinct epithelial features (C) and signaling pathways (D) in various epithelial cell subpopulations during PC progression.

E. Dot plot showing the expression of representative genes associated with indicated signaling pathways in various epithelial cell subpopulations during PC progression.

**Figure S8**

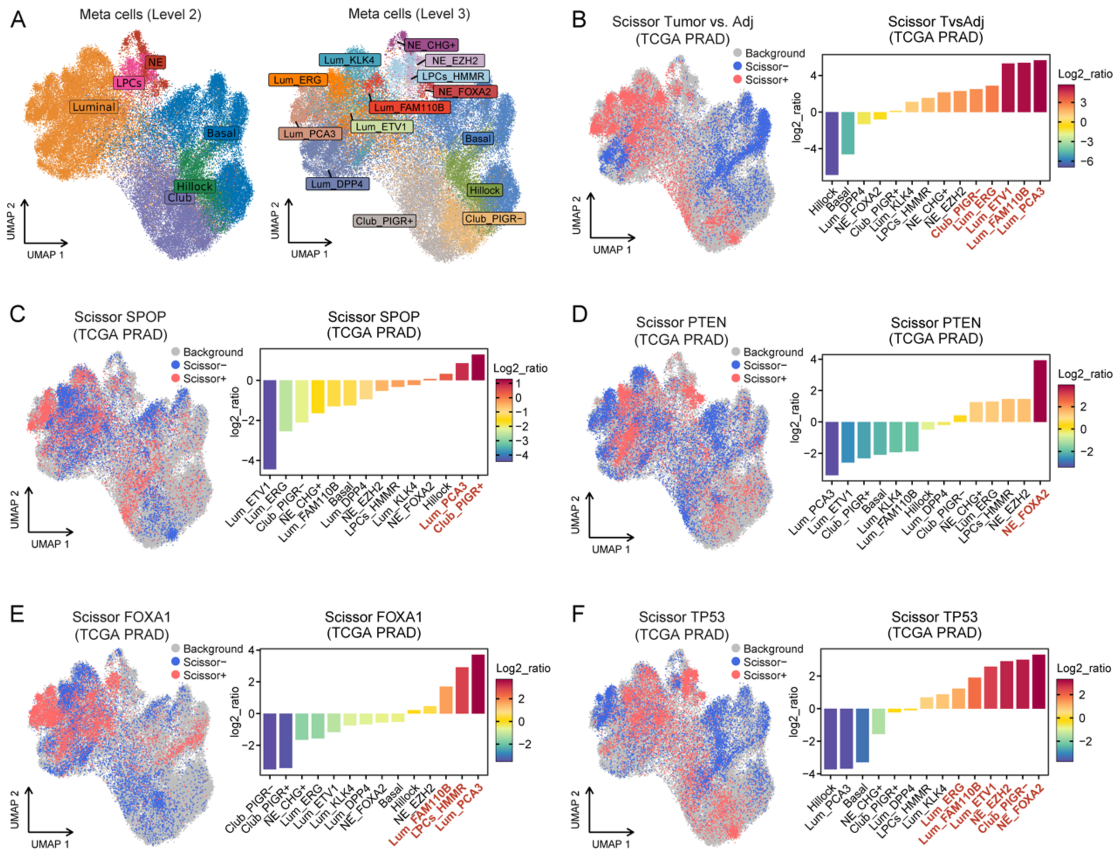


**Figure S8. Association of epithelial cellular composition and distinct phenotypes in the TCGA PRAD cohort, related to Figure 3.**

A. UMAP visualization of epithelial meta-cells, colored by meta-cell types (left) and meta-cell states (right). LPCs, lineage plasticity related cells; NE, neuroendocrine cells.

B-F. Scissor analysis relating phenotypic information from TCGA PRAD bulk RNA-seq data with epithelial cells, including PRI versus adjacent (B), SPOP mutation (C), PTEN mutation (D), FOXA1 mutation (E), and TP53 mutation (F). UMAP plots indicate the position of cells positively (red) or negatively (blue) associated with tumor or gene mutations. A log2 ratio >0 indicates a positive association with tumor or mutation, respectively.

**Figure S9**

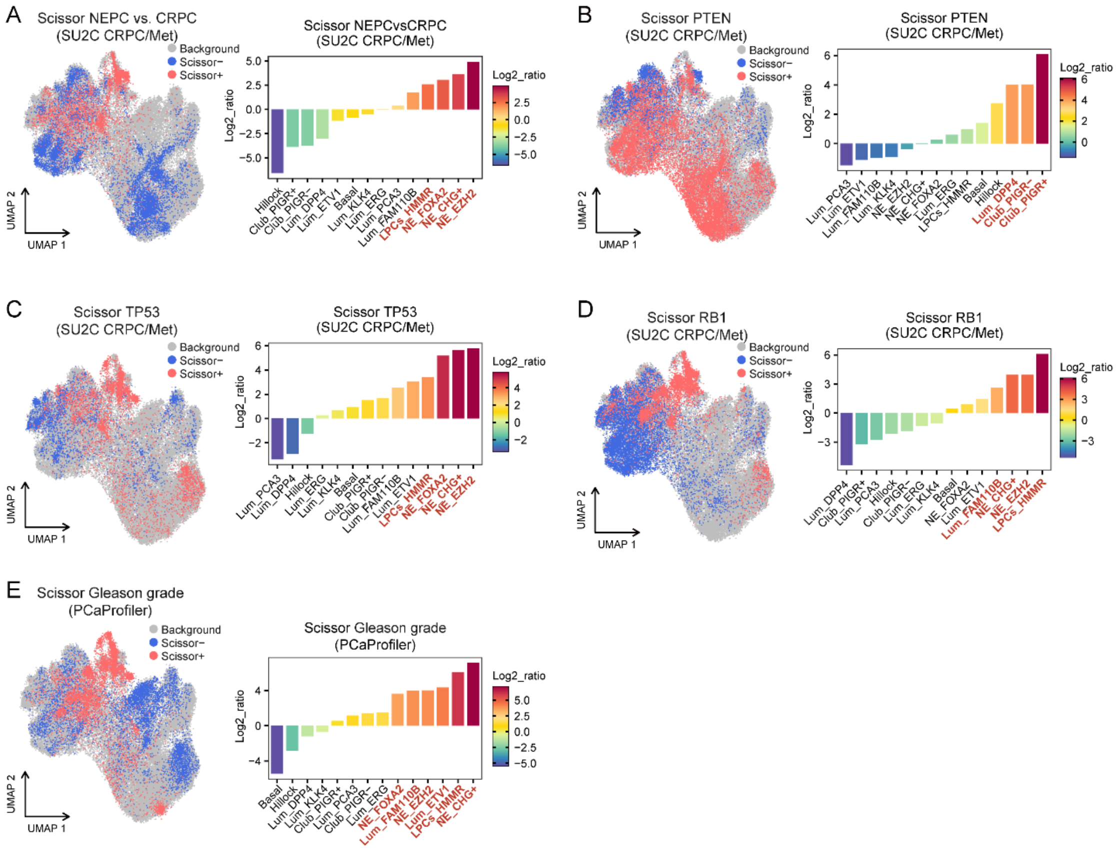


**Figure S9. Association of epithelial cellular composition and distinct phenotypes in the advanced PC datasets, related to Figure 3.**

A-D. Scissor analysis relating phenotypic information from SU2C CRPC/Met bulk RNA-seq data with epithelial meta-cells, including NEPC versus CRPC (A), PTEN mutation (B), TP53 mutation (C), and RB1 mutation (D).

E. Scissor analysis showing progression related epithelial cells using PCaProfiler bulk RNA-seq with epithelial meta-cells.

UMAP plots indicate the position of cells positively (red) or negatively (blue) associated with NEPC, mutation or Gleason grade. A log2 ratio >0 indicates a positive association with NEPC, mutation or Gleason grade, respectively.

#### Figure S10

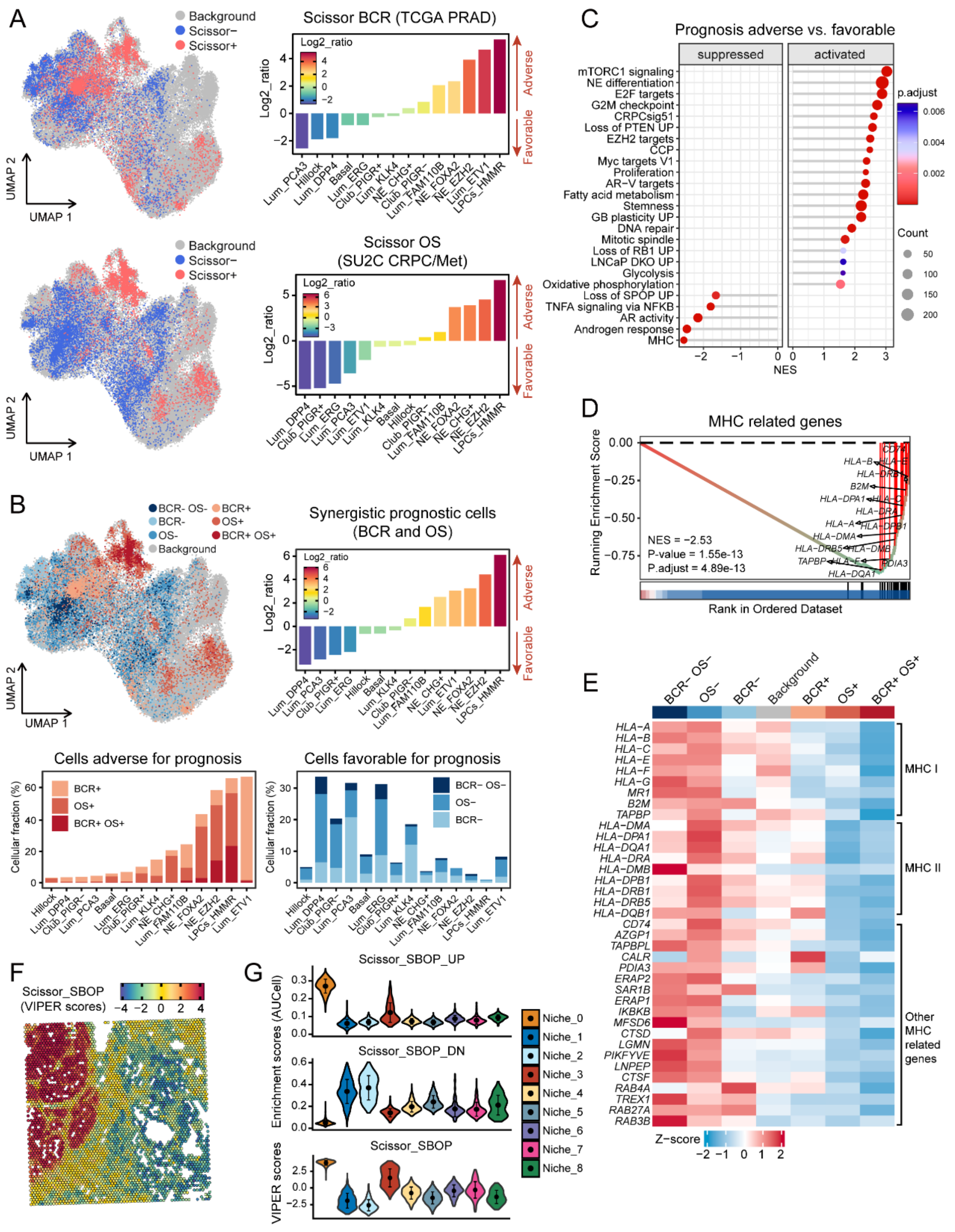


**Figure S10. Identification of key epithelial cell subpopulation and molecular characteristics linked to an adverse prognosis, related to Figure 3.**

A. Scissor analysis showing poor prognosis related epithelial cells based on BCR of TCGA PRAD dataset and OS of SU2C CRCP/Met dataset. UMAP plots indicate the position of cells positively (red) or negatively (blue) associated with poor prognosis. A log2 ratio >0 indicates a positive association with poor prognosis.

B. UMAP plot showing the distribution of synergistic prognostic cells by combining BCR and OS associated epithelial cells (left), and bar plot showing log2 ratio (upper) or cellular fraction (lower) of synergistic prognostic cells for each cell subpopulation.

C. GSEA showing the activated and suppressed pathways between predicted survival-adverse and favorable epithelial cells. NES, normalized enrichment score.

D. GSEA plots showing the suppressed activity of MHC signaling in survival-adverse epithelial cells versus survival-favorable epithelial cells.

E. Heat map showing the expression of representative genes associated with indicated signaling in different Scissor groups related to prognosis.

F. The spatial feature plot of the Scissor-derived gene signature associated with prognosis (SBOP).

G. Violin plots showing the spatial distribution of the SBOP scores among different spatial niches.

**Figure S11**

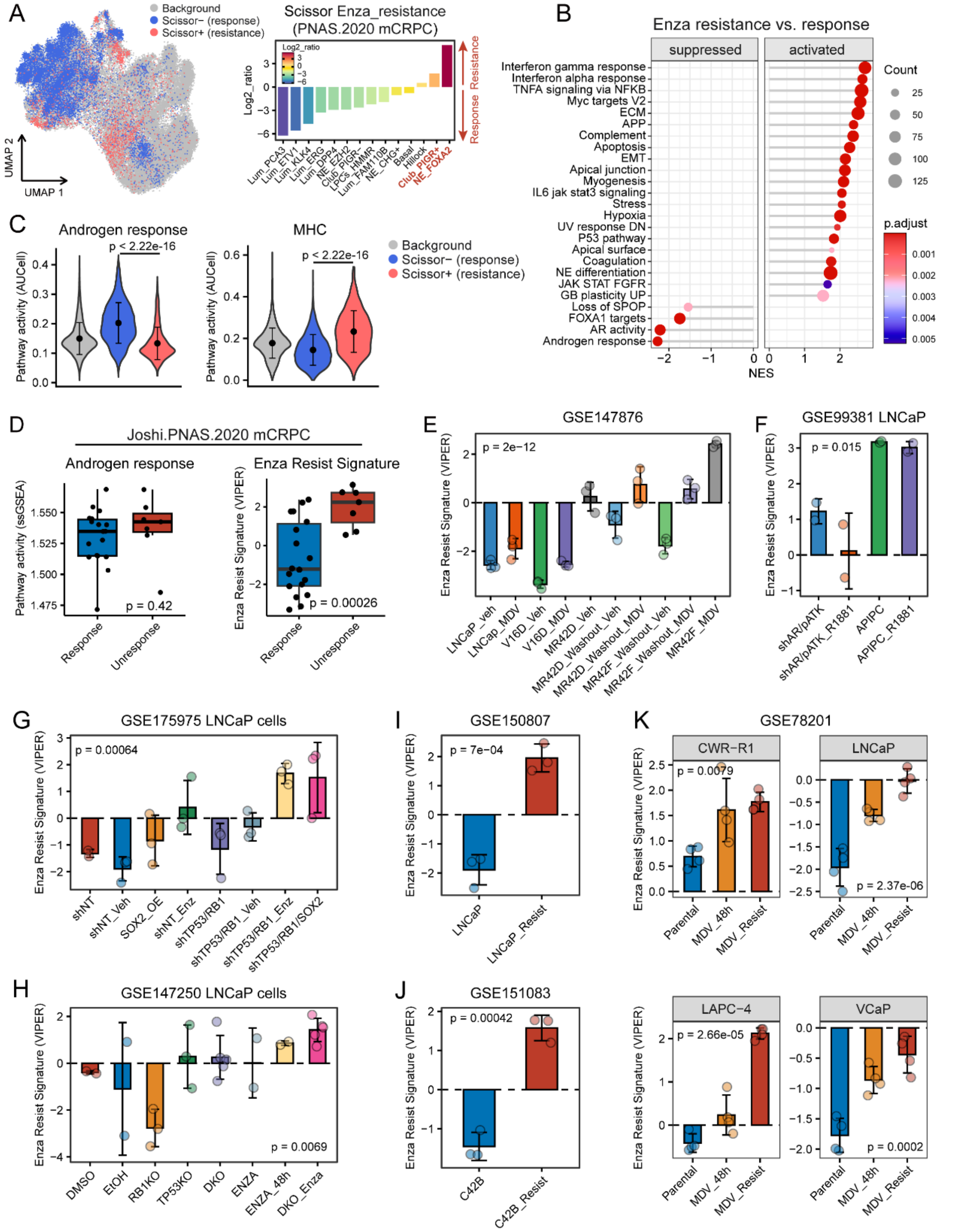


**Figure S11. Association of epithelial molecular characteristics and enzalutamide resistance in the advanced PC data, related to Figure 3.**

A. Scissor analysis showing enzalutamide resistance-related epithelial cells based on our in-house trial, included thirty-six patients with mCRPC who had not previously received enzalutamide. UMAP plots indicate the position of cells positively (red) or negatively (blue) associated with enzalutamide resistance. A log2 ratio >0 indicates a positive association with enzalutamide resistance.

B. Gene set enrichment analysis (GSEA) showing the activated and suppressed pathways between predicted enzalutamide resistance and response epithelial cells. NES, normalized enrichment score.

C. AUCell enrichment analysis comparing the activities of androgen response (left) and AR pathway (right) among different Scissor groups related enzalutamide resistance.

D. Bar plots showing the activity of the androgen response pathway (left) and Scissor-derived enzalutamide resistance signature (Enza Resist Signature, right) among patients with enzalutamide response and un-response.

E-K. Bar plots showing the activity of Enza Resist Signature among different enzalutamide resistance or sensitive cell lines from published or our in-housed datasets, including GSE147876 (E), GSE99381 (F), GSE175975 (G), GSE147250 (H), GSE150807 (I), GSE151083 (J), and GSE78201 (K). Veh, vehicle DMSO; MDV, Enzalutamide. shAR, doxycycline-inducible shRNA targeting the AR; pATK, androgen-driven thymidine kinase gene; ARIPC, AR program-independent PC. shNT, control non-targeting short hairpin RNA; SOX2-OE, SOX2 overexpression; shTP53/RB1, TP53/RB1 deficiency. DKO, the dual knockout of TP53 and RB1. Error bars denote SD (C-K). Wilcoxon rank sum test (C, D, I and J). Kruskal-Wallis test (E-H, and K).

**Figure S12**

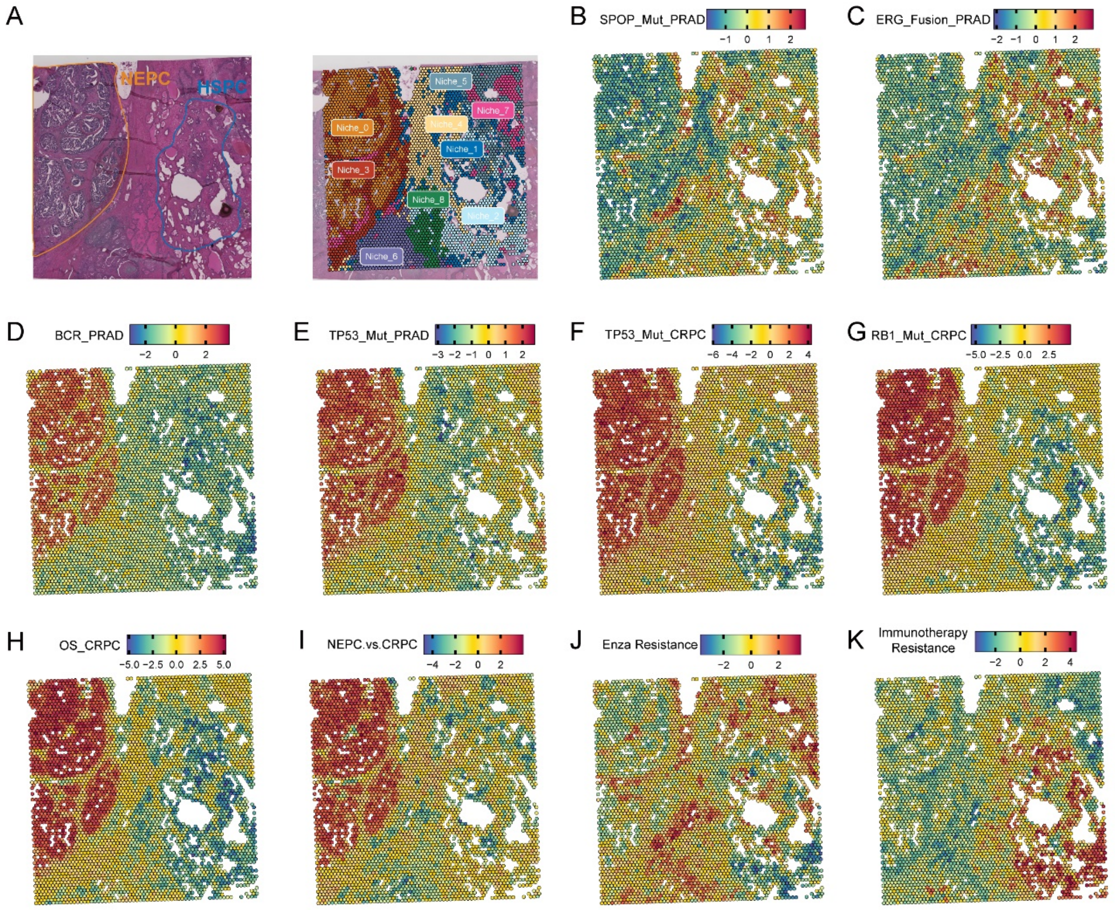


**Figure S12. The spatial distribution of genotype-phenotype based gene signature scores, related to Figure 3.**

A. The pathological annotation (left) and cell niches (right) of spatial RNA-seq data from the mixed section of NEPC and HSPC.

B-K. The spatial feature plot of the Scissor-derived gene signatures associated with SPOP mutation (B), ERG fusion (C), BCR (D), and TP53 mutation in TCGA PRAD data, as well as TP53 mutation (F), RB1 mutation (G), OS (H), NEPC (I), enzalutamide resistance (J) and immunotherapy resistance (K) in the advanced PC datasets.

**Figure S13**

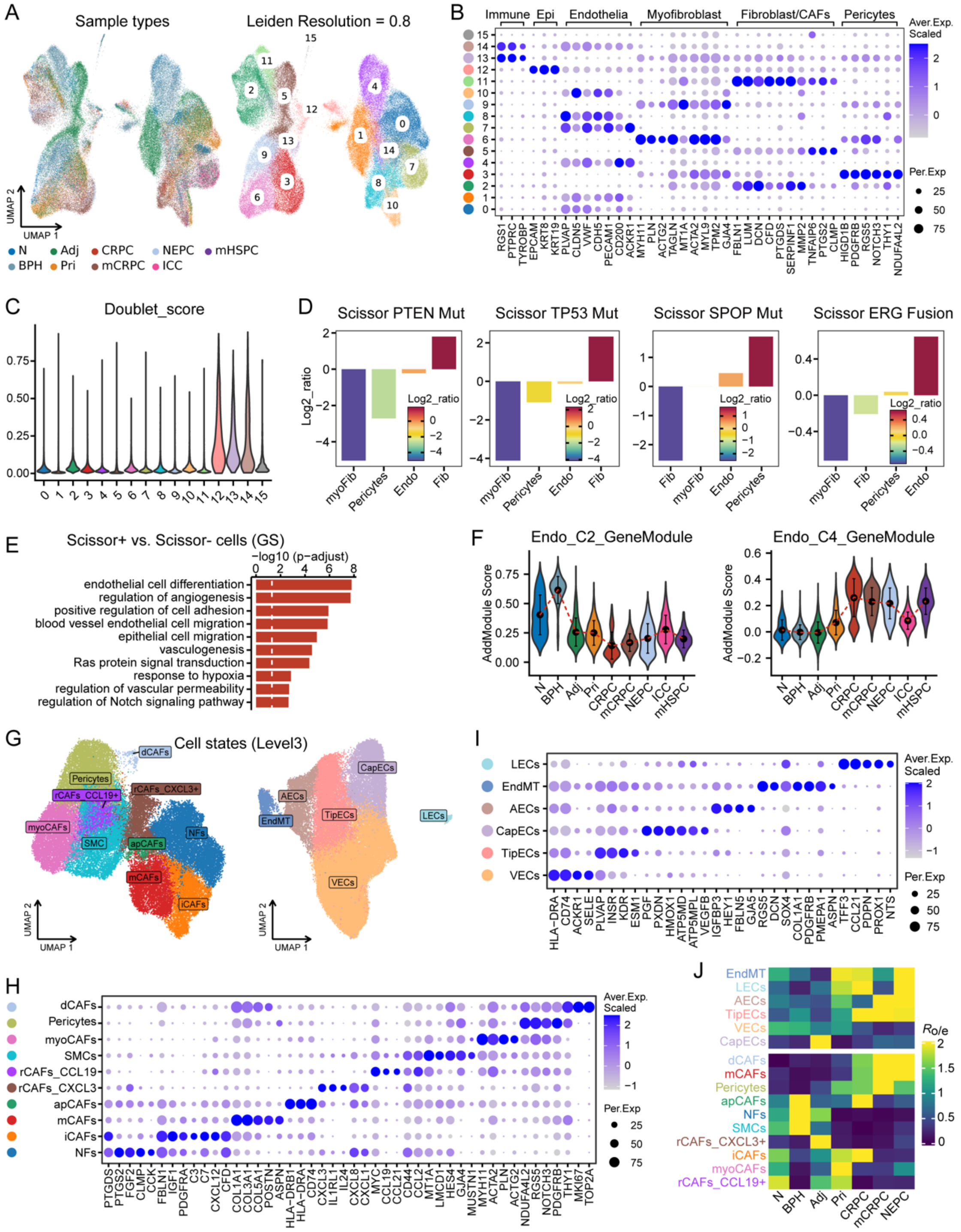


**Figure S13. Re-cluster analysis of stromal cell, related to Figure 4.**

A. UMAP plots showing 112,249 single stromal cells that colored by different disease groups (left) and cell clusters (right).

B. Dot plot of representative marker genes for each cluster.

C. Violin plots showing the doublet scores inferred by Scrublet for each cell cluster.

D. Bar plots showing phenotype related stromal cells identified by Scissor with TCGA PRAD bulk RNA-seq data, including PTEN mutation (A), TP53 mutation (B), SPOP mutation (C), and ERG fusion (D). A log2 ratio >0 indicates a positive association with gene mutations or ERG fusion, respectively.

E. Heat map showing normalized Jaccard similarity scores between UP or DN Scissor DEGs and different gene clusters generated from Mfuzz time-series analysis. UP, up-regulated expression in Scissor+ cells. DN, down-regulated expression in Scissor+ cells.

F. GO enrichment analysis of the up-regulated DEGs in Scissor+ cells versus Scissor- cells derived by Gleason grade.

G. UMAP plots depicting 10 fibroblast cell subclusters (left) and 5 endothelial cell subclusters (right).

H and I. Dot plots of representative marker genes for each fibroblast (I) or endothelial (J) cell subcluster.

J. Tissue preference of each stromal cell subcluster estimated by Ro/e score.

**Figure S14**

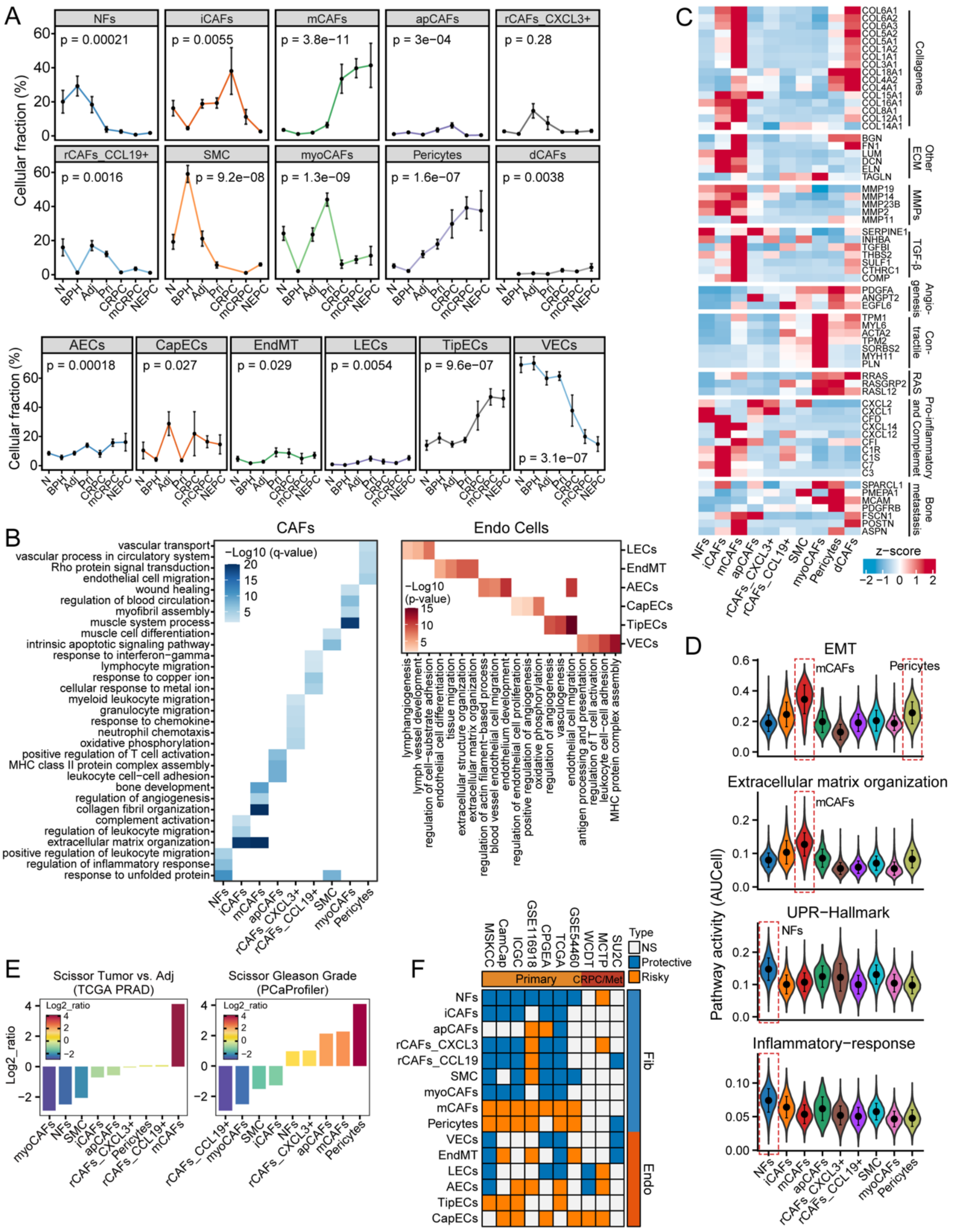


**Figure S14. Changes in cell proportion and molecular characteristics of stromal cells during the PC progression, related to Figure 4.**

A. Dynamic changes in cellular fraction for each stromal cell clusters during PC progression.

B. GO enrichment analysis of the top gene markers for each fibroblast (left) or endothelial (right) cell subcluster.

C. Heat map showing the expression of representative genes associated with indicated signaling in different fibroblast cell subclusters.

D. AUCell enrichment analysis showing distinct pathway characteristics in various fibroblast cell subclusters.

E. Bar plots showing phenotype related fibroblast cells identified by Scissor, including tumor vs. Adj (left) and Gleason grade (right). A log2 ratio >0 indicates a positive association with tumor or Gleason grade, respectively.

F. Univariate Cox regression analysis for each stromal subcluster. Survival index using BCR for primary PC and OS for CRPC/Met PC.

#### Figure S15

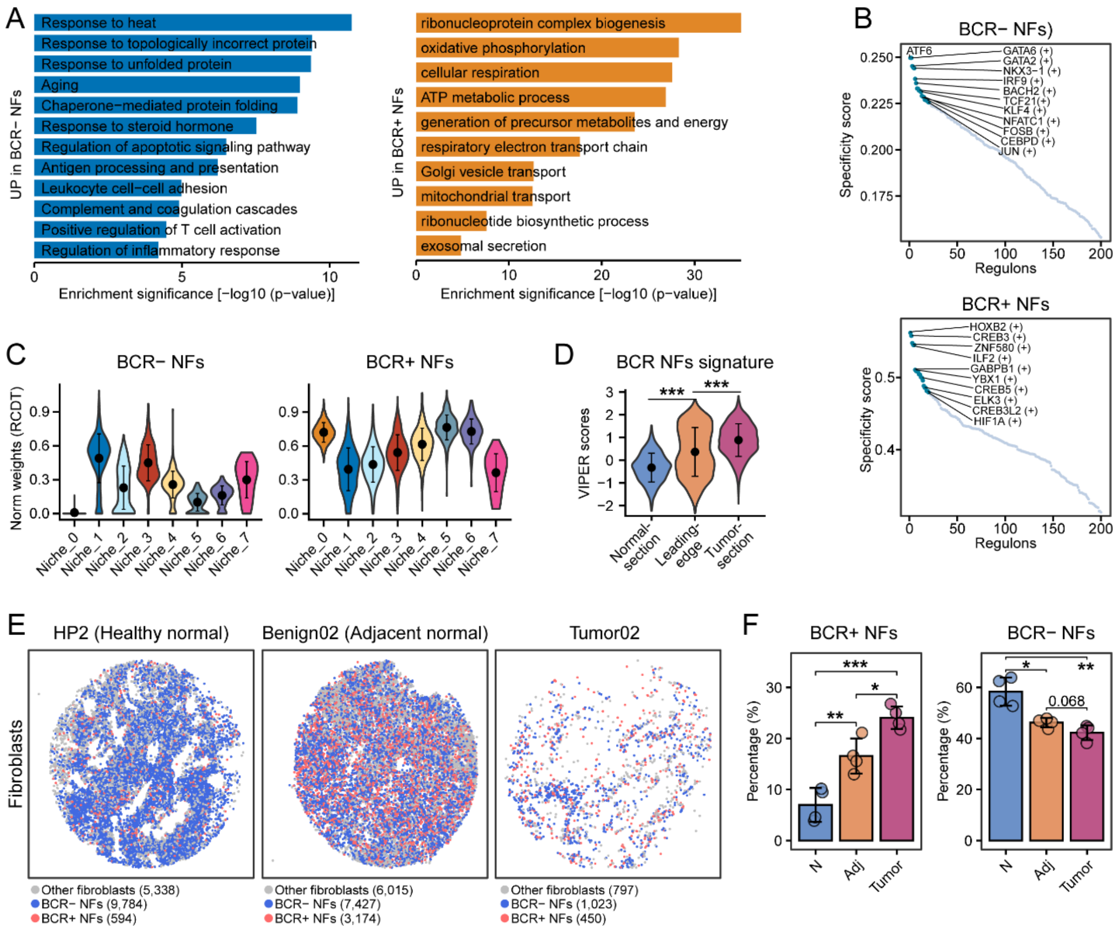


**Figure S15. The molecular characteristics and spatial distribution of BCR+ like NFs of PC, related to Figure 4.**

A. GO enrichment analysis of the up-regulated DEGs in BCR- like normal fibroblast (NF) cells (left) or BCR+ like NF cells (right). Scissor-related NFs were identified by combining all fibroblast cells with BCR in TCGA PRAD bulk RNA-seq data using the Scissor algorithm in COX-mode.

B. Scatter plot showing the specificity scores of regulons of BCR- like NF cells (upper) or BCR+ like NF cells (lower). The top 10 regulons are highlighted.

C. Violin plots showing the distribution of normalized weights for BCR- like NF cells (left) and BCR+ like NF cells (right) among different PC niches in the invasion leading edge section. Normalized weights were inferred by RCDT algorithm.

D. The activity of the Scissor-derived gene signature (BCR NFs signature), based on DEGs of

BCR+ like NF cells versus BCR- like NF cells, varies across the normal prostate section, the invasion leading edge section, and the tumor core section.

E. The spatial distribution of BCR-related NFs among normal, adjacent and tumor sections generated from Slide-seq V2 technology. Cells were inferred by RCTD algorithm.

F. Quantitative analyses of the percentage of BCR+ like NFs (right) or BCR- like NFs (right) in fibroblast cells for each disease group (sample = 3). Each dot represents an individual sample. Error bars denote SD (C, D and F). Kruskal-Wallis test with Bonferroni correction (D and F). *, $p < 0.05$; **, $p < 0.01$; ***, $p < 0.001$.

**Figure S16**

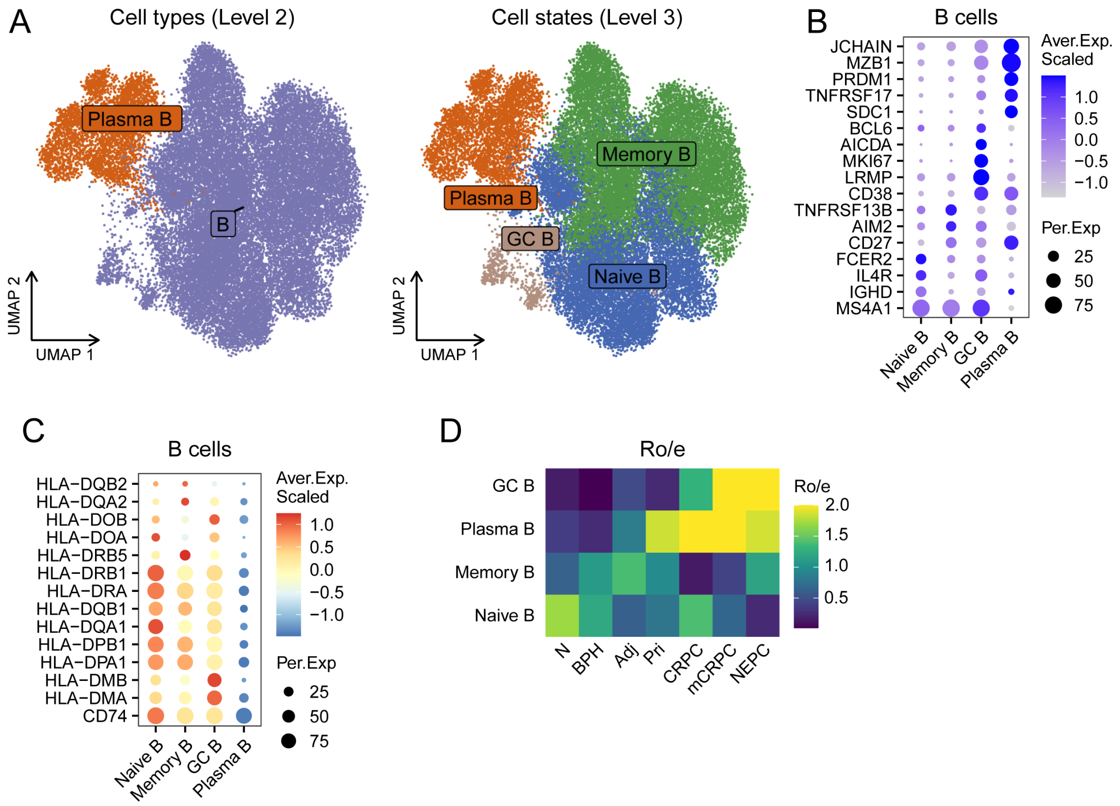


**Figure S16. Characteristics of B cells during PC progression, related to Figure 5.**

A. UMAP plots showing a total of 30,860 B cells, colored by cell types (level2, left) and cell states (level3, right). GC B, germinal center B cells.

B. Dot plot of representative marker genes for each B cell cluster.

C. Dot plot showing the expression levels of MHC-II genes for B cell clusters.

D. Tissue preference of each B cell cluster estimated by Ro/e score.

**Figure S17**

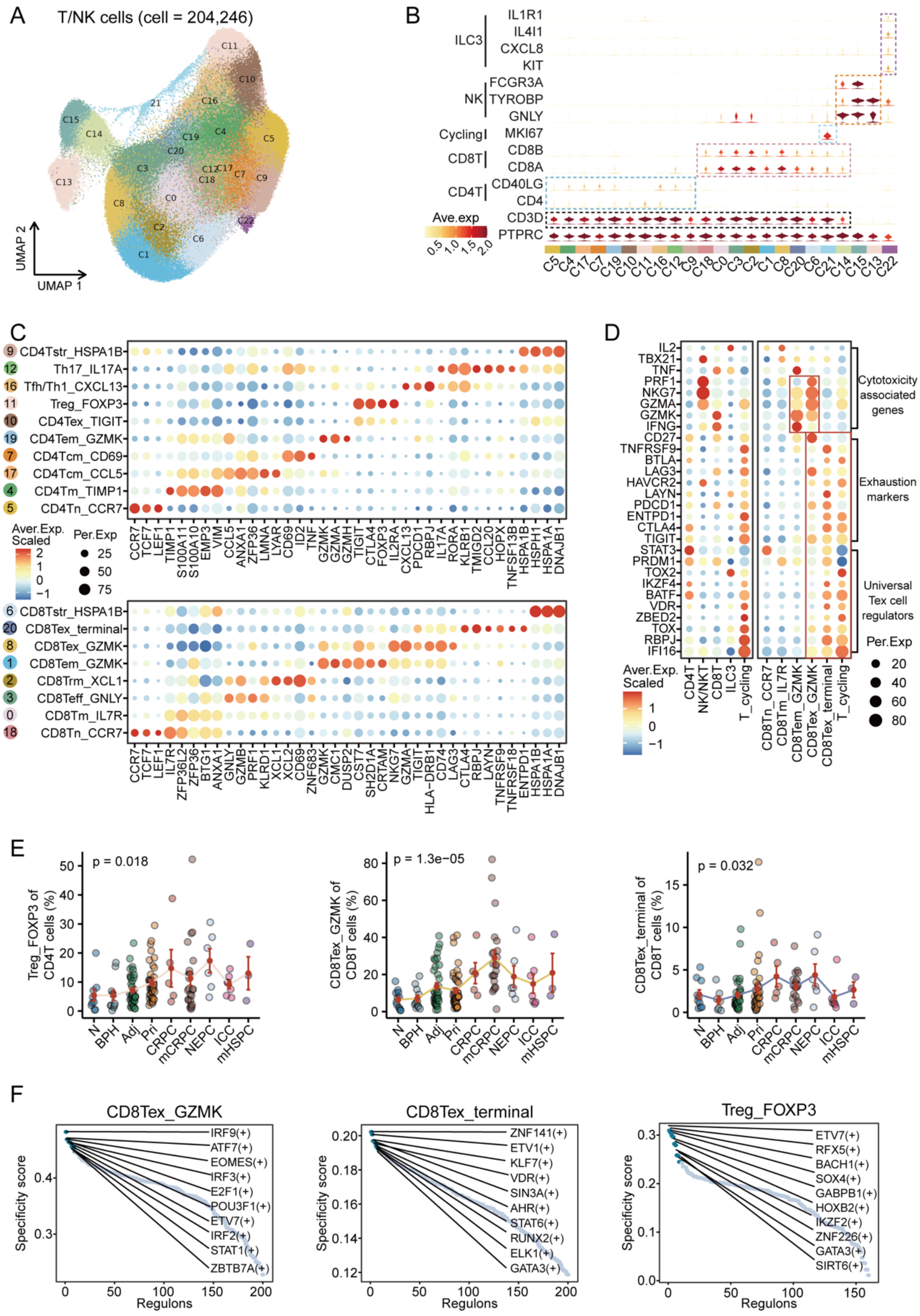


**Figure S17. Characteristics of T cells during PC progression, related to Figure 5.**

A. UMAP plots showing 204,246 single T/NK cells that colored by different cell clusters.

B. Violin plot of representative marker genes for each cluster.

C. Dot plot showing the expression of representative marker genes for each CD4+ T (upper) and CD8+ T (lower) cell subpopulations.

D. Dot plot showing the expression of representative genes associated with indicated signaling among different cell types (left) and cell states (right).

E. Scatter diagram showing the dynamic changes in frequency of Treg_FOXP3 (left), CD8Tex_GZMK (middle) and CD8Tex_terminal (right) cells with PC progression.

F. Scatter plot showing the specificity scores of regulons of Treg_FOXP3 (left), CD8Tex_GZMK (middle) and CD8Tex_terminal (right) cells. The top 10 regulons are highlighted.

**Figure S18**

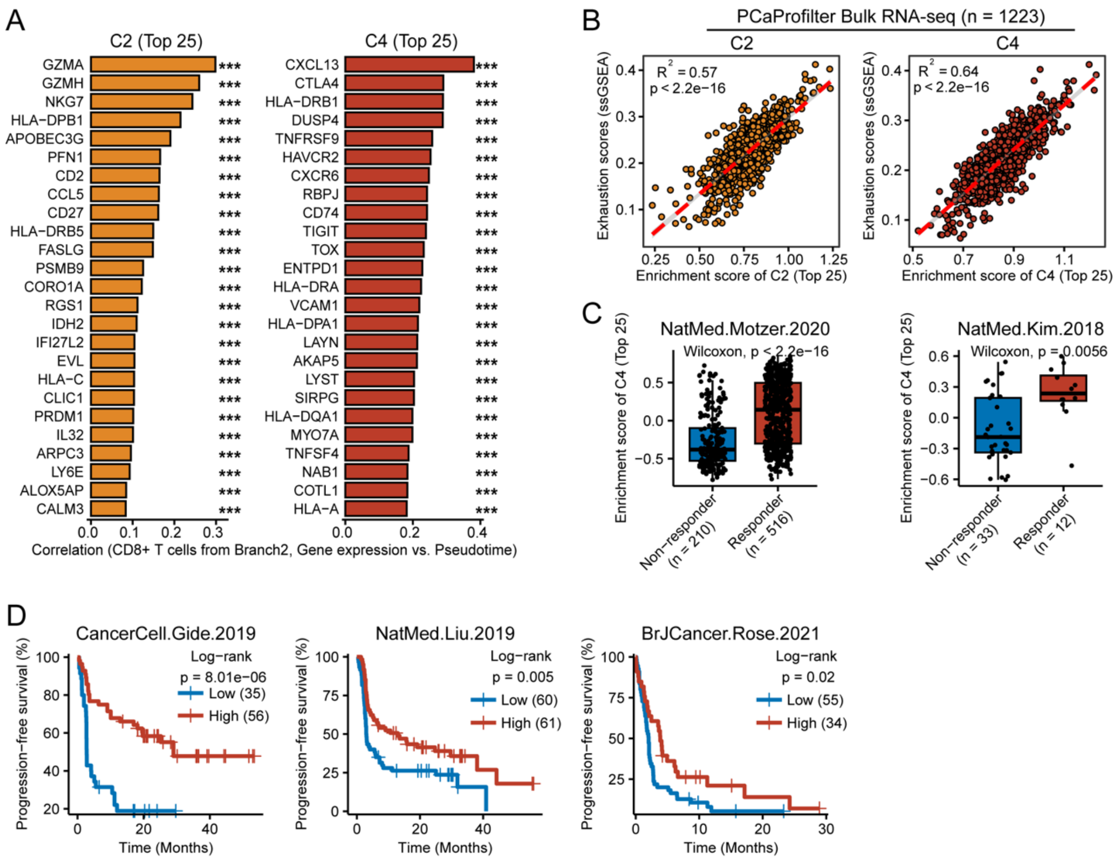


**Figure S18. Characteristics of T cells during PC progression, related to Figure 5.**

A. Bar plots showing top 25 genes from C2 (left) and C4 (right), whose gene expression was positively correlated with cell pseudotime in CD8T+ cells from branch 2.

B. Scatter diagram showing the correlation between gene signature scores and exhausted T cell signature scores in PCaProfilter bulk RNA-seq dataset. Each dot represents an individual sample.

C. The distribution of the C4 gene cluster signature among patients with immunotherapy response and non-response in two published cohorts.

D. Kaplan–Meier survival analysis illustrating overall survival outcomes for patient groups with low and high scores of the C4 gene cluster signature in three immunotherapy cohorts.

**Figure S19**

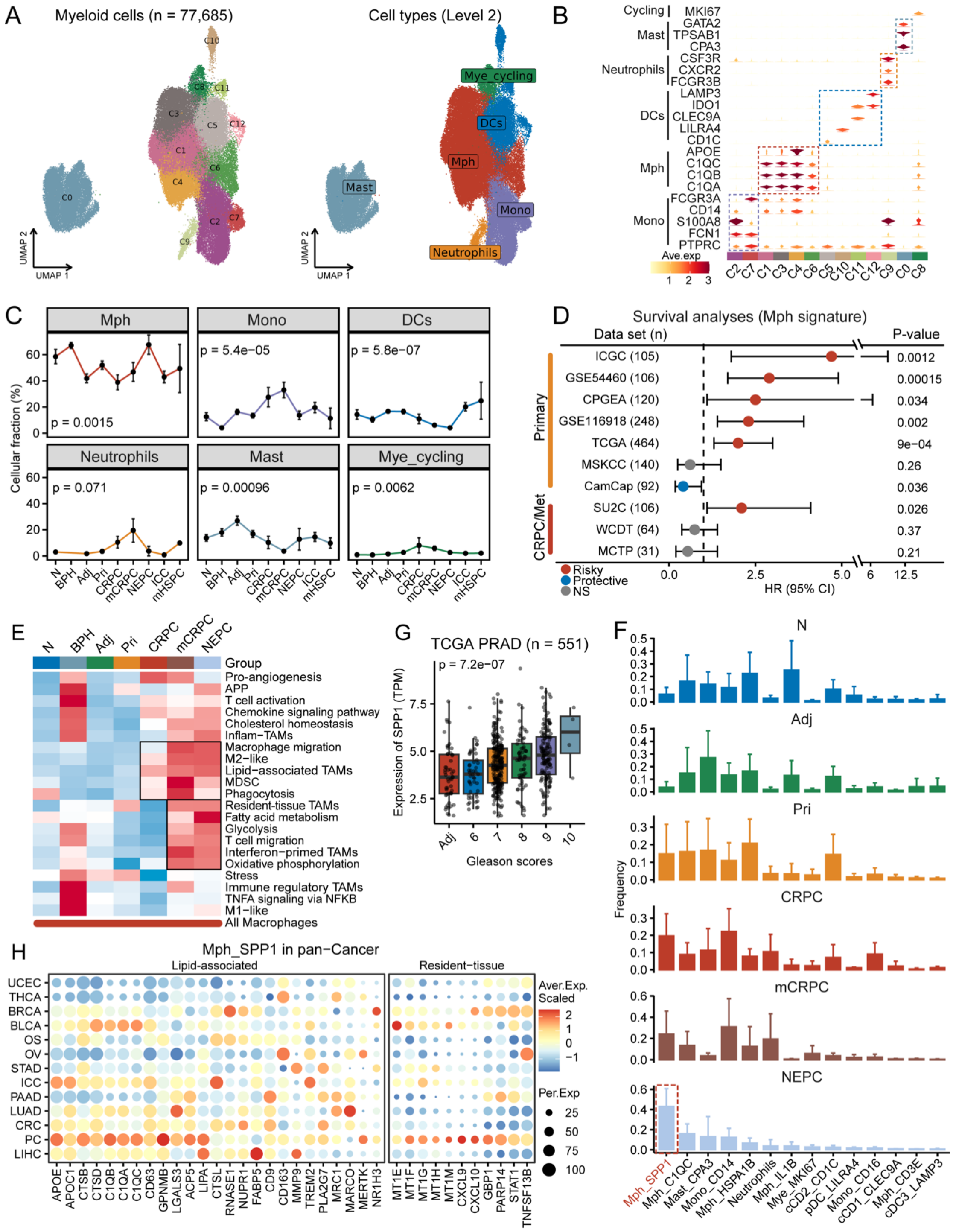


**Figure S19. Characteristics of macrophage cells during PC progression, related to Figure 6.**

A. UMAP plots showing 77,685 single myeloid cells that colored by different cell clusters (left) and cell types (right).

B. Violin plot of representative marker genes for each cluster.

C. Dynamic changes in cellular fraction for each myeloid cell types during PC progression.

D. Forest plot illustrating survival analysis of the macrophage gene signature. HR, Hazard ratio; Error bars denote 95% confidence interval (CI).

E. Heatmap illustrating dynamic changes in activities of 21 curated gene signatures during PC progression.

F. Bar plots showing the myeloid cell compositions in the normal, adjacent, PRI, CRPC, mCRPC and NEPC groups.

G. Box plot showing the dynamic changes in expression of SPP1 with PC progression.

H. Dot plot showing the expression of representative genes associated with indicated pathways in Mph_SPP1 cells from different cancer types.

### Figure S20

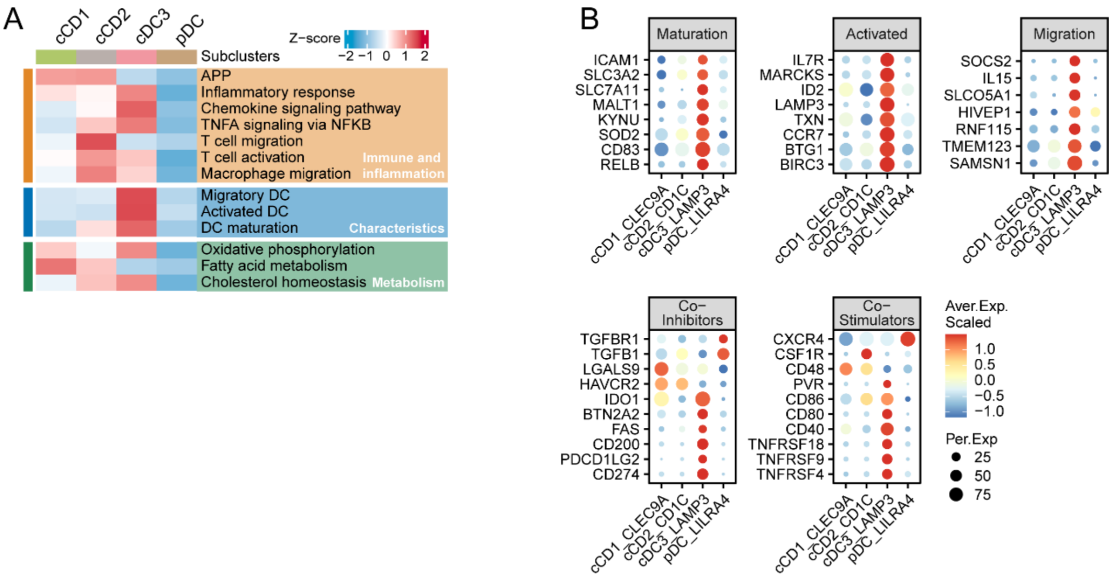


**Figure S20. Characteristics of DC cells during PC progression, related to Figure 6.**

A. Heatmap showing the activities of 13 curated gene signature for each DC cell subcluster.

B. Dot plot showing the expression of representative genes associated with indicated signaling in DC cell subclusters.

**Figure S21**

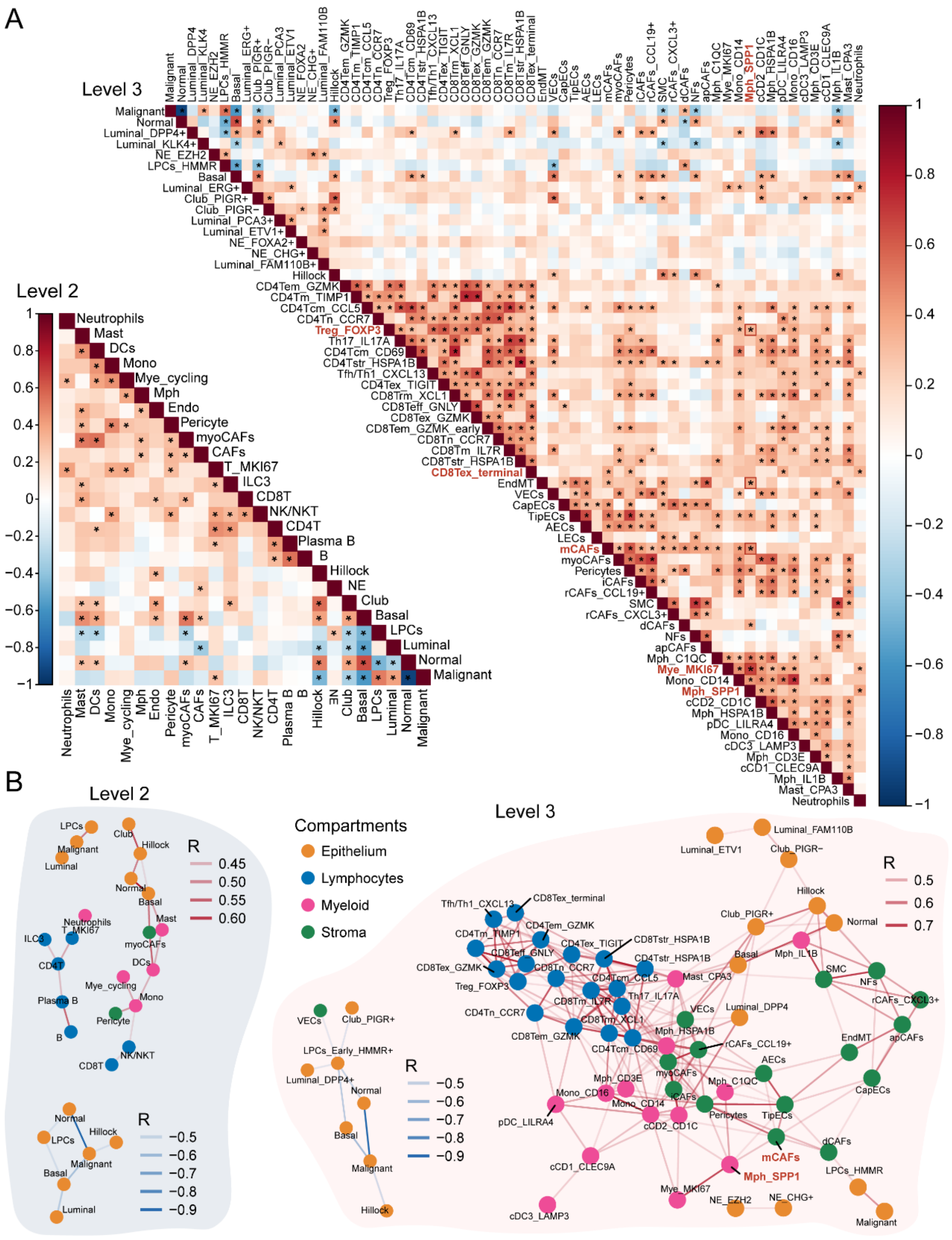


**Figure S21. Subpopulation proportion correlation network of PC TME, related to Figure 7.**

A. Heatmap represents the correlation matrix across subpopulation proportions. Colored row and column tracks represent the cellular compartments used to derive % calculations. *, adjust p-value $< 0.05$.

B. Positively correlated ($|R| > 0.4$) feature nodes visualized on a correlation-based network layout. Node colors represent the derived compartments (legend). Opacity of connections (edges) represent Spearman correlation R value. Blue line represents negative correlation, and red line represents positive correlation.

**Figure S22**

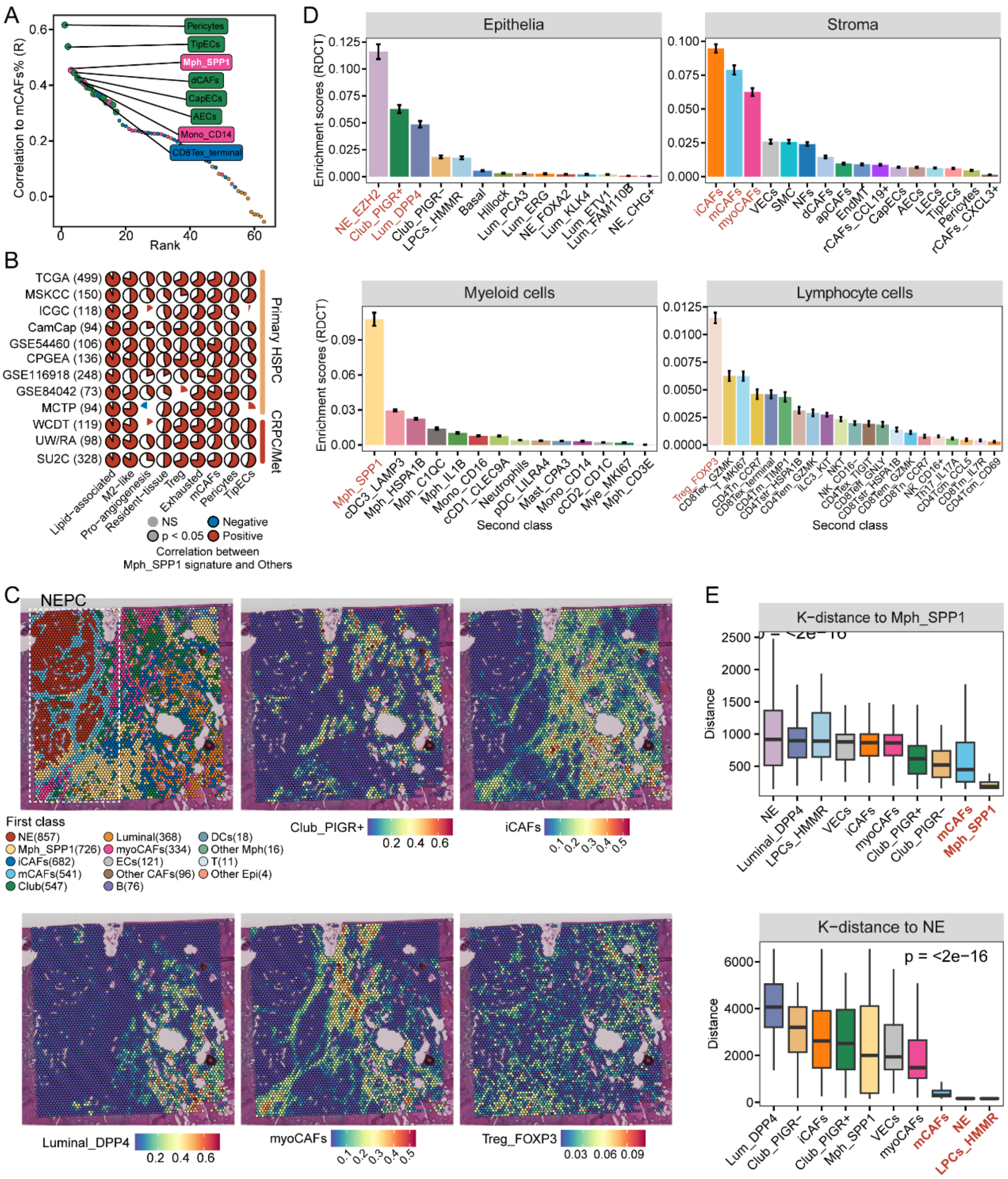


**Figure S22. Spatial distribution and abundance of PC cell subpopulation, related to Figure 7.**

A. Spearman correlation analyses between the cell proportion of mCAFs and other cell subpopulations. Each data point on the scatter plot represents a cell subpopulation, ordered by its correlation to mCAFs.

B. Correlation analysis of Mph_SPP1 gene signature scores with activities of multiple signaling pathways or cell type signatures in 12 bulk transcriptomic cohorts.

C. The rough define cell types (first class) of each spots (left), as well as the spatial distribution of Club_PIGR+ (middle) and iCAFs (right), as inferred by the RCTD deconvolution algorithm.

D. Enrichment scores of cell subpopulations (second class) for each cellular compartment.

E. Box plot showing RCTD-based spatial K-distance of Mph_SPP1 (upper) or NE tumor cells (lower) to other cell subpopulations.

#### Figure S23

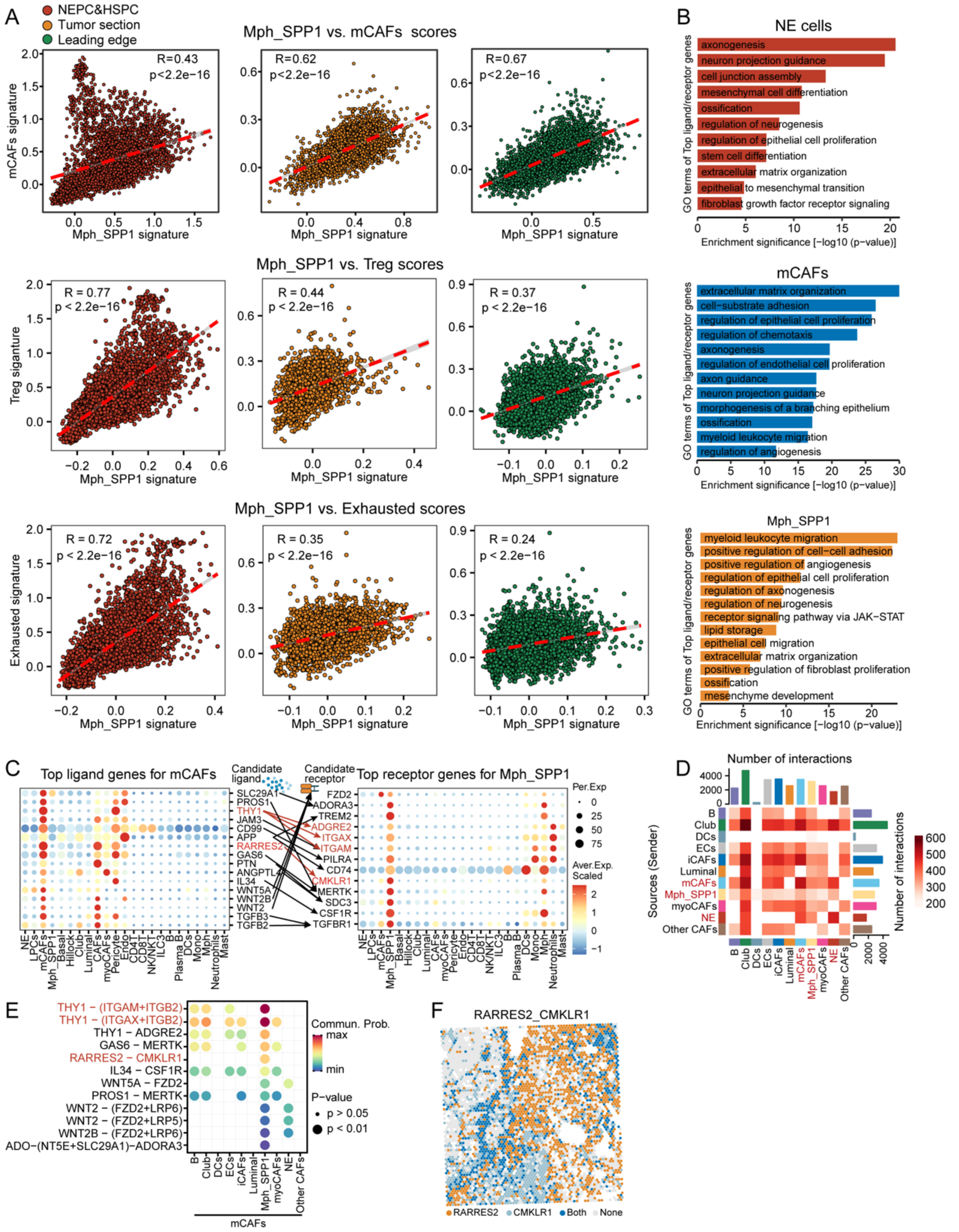


**Figure S23. Spatial distribution and abundance of PC cell subpopulation, related to Figure 7.**

A. Spearman correlation analyses between the cell proportion of mCAFs and other cell subpopulations. Each data point on the scatter plot represents a cell subpopulation, ordered by its correlation to mCAFs.

B. Correlation analysis of Mph_SPP1 gene signature scores with activities of multiple signaling pathways or cell type signatures in 12 bulk transcriptomic cohorts.

C. The rough define cell types (first class) of each spots (left), as well as the spatial distribution of Club_PIGR+ (middle) and iCAFs (right), as inferred by the RCTD deconvolution algorithm.

D. Enrichment scores of cell subpopulations (second class) for each cellular compartment.

E. Box plot showing RCTD-based spatial K-distance of Mph_SPP1 (upper) or NE tumor cells (lower) to other cell subpopulations.

F. The feature plots showing the expression distribution of ligand-receptor gene pair by binarizing their expression in each spot. Each spot is colored based on whether it only expresses the ligand, the receptor or both signaling molecules

#### Figure S24

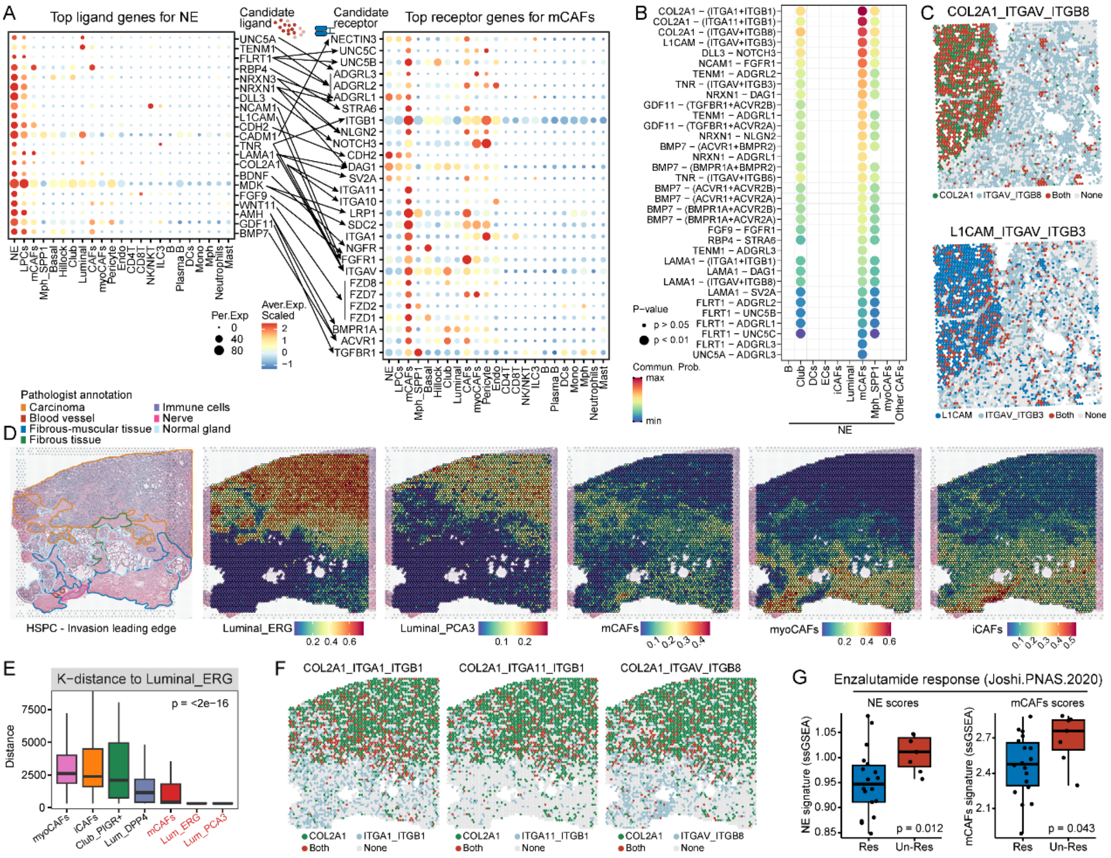


**Figure S24. Identifying potential ligand pairs involving interactions between mCAFs and Mph_SPP1, related to Figure 7.**

A. Dot plots showing the expression of candidate ligand-receptor gene pairs involved the interaction between NE tumor cells and mCAFs.

B. All the significant ligand-receptor pairs that contribute to the interactions from mCAFs to other subclusters in spatial tissue section. Dot size indicated the statistical significances by permutation test. Dot color indicated the mean interaction strength levels.

C. The feature plots showing the expression distribution of each ligand-receptor gene pair by binarizing their expression in each spot of NEPC&HSPC tissue section. Each spot is colored based on whether it only expresses the ligand, the receptor or both signaling molecules.

D. The pathological annotation (left) ), as well as the spatial distribution of each high enriched cell subpopulations in the invasion leading edge spatial section of primary PC.

E. Box plot showing RCTD-based spatial K-distance of Luminal_EGR+ cells to other cell subpopulations.

F. The feature plots showing the expression distribution of each ligand-receptor gene pair by binarizing their expression in each spot of the invasion leading edge tissue section. Each spot is colored based on whether it only expresses the ligand, the receptor or both signaling molecules.

G. The distribution of NE cell gene signature scores and mCAFs gene signature scores between enzalutamide response groups in our in-house cohort.

#### Supplementary Methods

##### RCTD deconvolution analysis

The RCTD method[1] was employed to deconvolute each spot in the 10X Visium or Slide-seqV2 slices using scRNA-seq data as a reference. RCTD objects were created for each slice from the processed RDS objects, with parameter 'max_cores = 4' and 'test_mode = FALSE'. RCTD pipeline was run on the RCTD objects, with parameter 'doublet_mode = full' for 10X Visium data or 'doublet_mode = doublet' for Slide-seq V2 data. The deconvolution results were matrices of celltype weights for each spot. The cell type weights were normalized to make the sum of cell type weights in each spot equal to 1. Proportion matrices, whose rows indicate spots and columns indicate cell types (consider it a second class), were then created and stored in the analyzed spatial transcriptome for loading. For the rough annotation, we assigned each spot to a specific cell type as the first class based on the highest probabilistic proportion.

##### SuperCell algorithm for calculating metacells

To succinctly describe the observed phenotypic diversity, we employed R package SuperCell[2] to compute metacells separately for each sample and cell subcluster using the SCimplify function with parameter 'gamma=20', 'n.var.genes=1000', 'k.knn=5', 'n.pc=10', 'cell.annotation=ct.L3' and 'cell.split.condition=SampleID'. The metacell approach groups similar single cells from each sample into distinct, highly detailed cell states. This ensures that any variations among cells within a metacell are more likely due to technical factors rather than biological differences effectively reducing noise and addressing the sparsity inherent in scRNA-seq data[2]. By aggregating counts within each metacell, we obtained robust and comprehensive transcriptomic quantification for each cell state, effectively reducing noise and addressing the sparsity inherent in scRNA-seq data. Furthermore, metacells offer the advantage of enumerating observed cell states that are more amenable to comparative analyses.

##### Scissor analysis

Using our Scissor algorithm[3] with cutoff set as 0.2, we investigated phenotype associated cell subclusters and molecular characteristics at single-cell resolution by combining transcription expression datasets and our PCCAT. For binary categorical phenotypes, logistic regression model was applied. For BCR or OS survival data, cox-regression model was applied. For continuous phenotype Gleason grade in PCaProfiler bulk dataset, linear regression model was applied. Furthermore, we calculated the proportion of Scissor+ cells (positively correlated with worse survival, mutation, treatment non-response, and tumor progression), Scissor- cells (negatively correlated), and the log2 ratio between the proportions of Scissor+ and Scissor- cells. Differential expression analysis was performed to compare the expression profile of Scissor+ and Scissor- cells. We defined DEGs based on a cutoff of $|avg\_log2FC| > 0.8$ and an adjusted p-value (p_val_adj) of $< 0.01$. Then Scissor scores were calculated using the Virtual Inference of Protein activity by Enriched Regulon analysis (VIPER) algorithm[4], which combined both the UP and DN Scissor-related signatures.

**Cell communication analysis**

To explore cell-to-cell communication patterns, we employed the CellChat2 method (v2.0.0)[5], following the official workflow. We assessed potential ligand/receptor interactions across all spots, with particular focus on interactions among Mph_SPP1, mCAFs and NE cells. This analysis was conducted through functions such as ‘computeCommunProb’, ‘computeCommunProbPathway’, ‘aggregateNet’, and ‘spatialFeaturePlot’, all using standard parameters.

**Streamlined single-cell PC classifier**

Leveraging the Python package “Celltypist” (v.1.5.0)[6], a single-cell label transfer framework, we trained a logistic regression with stochastic gradient descent learning model on the SuperCell atlas. Default parameters were used with the sole following exception (feature_selection = True). The resulting model was subsequently validated on two independent scRNA-seq datasets of human PC from recently published studies, achieving high precision and F1-scores: Masetti et al.[7] (Precision = 0.99, and F1-score = 0.99), and Hawley et al.[8]

(Precision = 0.99, and F1-score = 0.98).
